# High dimensional immunotyping of the obese tumor microenvironment reveals model specific adaptation

**DOI:** 10.1101/2020.10.14.338806

**Authors:** Cara E Wogsland, Hilde E Lien, Line Pedersen, Pahul Hanjra, Sturla M Grondal, Rolf A Brekken, James B Lorens, Nils Halberg

**Affiliations:** Department of Biomedicine, University of Bergen, N-5020 Bergen, Norway; Division of Surgical Oncology, Department of Surgery, and Hamon Center for Therapeutic Oncology Research; Department of Pharmacology, University of Texas Southwestern Medical Center, Dallas, Texas, USA

**Author notes:** **Corresponding Author:** Nils Halberg, Department of Biomedicine, University of Bergen, Jonas Lies vei 91, 5020 Bergen, Norway.

**Keywords:** tumor immunology, suspension mass cytometry, batch correction, obesity, breast cancer, pancreatic cancer

## Abstract

Obesity is a disease characterized by chronic low-grade systemic inflammation and has been causally linked to the development of 13 cancer types. Several studies have been undertaken to determine if tumors evolving in obese environments adapt differential interactions with immune cells and if this can be connected to disease outcome. Most of these studies have been limited to single cell lines and tumor models and analysis of limited immune cell populations. Given the multicellular complexity of the immune system and its dysregulation in obesity, we applied high-dimensional suspension mass cytometry to investigate how obesity affects tumor immunity. We used a 36-marker immune-focused mass cytometry panel to interrogate the immune landscape of orthotopic syngeneic mouse models of pancreatic and breast cancer. Unanchored batch correction was implemented to enable simultaneous analysis of tumor cohorts to uncover the immunotypes of each cancer model and reveal remarkably model-specific immune regulation. In the E0771 breast cancer model, we demonstrate an important link to obesity with an increase in two T cell suppressive cell types and a decrease in CD8 T-cells.

## Introduction

Obesity is a risk factor for at least 13 types of cancer including breast and pancreatic cancer (Genkinger et al, 2011; Picon-Ruiz et al, 2017; Pierobon & Frankenfeld, 2013; Renehan et al, 2015). In addition to being associated with risk of cancer, obesity correlates with worse prognosis and higher mortality rates among breast and pancreatic cancer patients (Calle et al, 2003; Chan & Norat, 2015; Chan et al, 2014; Choi et al, 2016; Yuan et al, 2013). The mechanisms by which obesity contributes to cancer development and outcome are currently incompletely understood (Donohoe et al, 2017). Obesity leads to local and systemic inflammation and immune system dysregulation characterized by increased levels of pro-inflammatory cytokines and pro-inflammatory and immunosuppressive immune cells (Apostolopoulos et al, 2016). Specific obesity-induced changes include increased abundance of immunosuppressive myeloid derived suppressor cells (MDSC), pro-inflammatory M1 and metabolically activated (MMe) macrophages, and associated crown-like structures found in obese adipose tissue (Coats et al, 2017; Hale et al, 2015; Kratz et al, 2014; Pawelec et al, 2019; Tiwari et al, 2019; Weisberg et al, 2003; Xu et al, 2003). Breast and pancreatic tumors are likely impacted by the local and systemic effects of obesity since these tumors develop in close proximity to mammary and omental adipose tissue, respectively.

Immunocompetent models of obesity and cancer are necessary to study immune changes in cancer in the obese environment. The C57Bl/6 mouse strain has been shown to have an obese phenotype when fed a high fat diet (HFD) including increased body mass, elevated blood sugar levels, and insulin resistance (Coats et al, 2017; Tiwari et al, 2019). When paired with such diet-induced obesity (DIO), syngeneic tumor cell lines can be used to study cancer associated immune system changes during obesity. Increased tumor incidence and accelerated tumor growth have been demonstrated in multiple obese murine models (Chung et al, 2020; Cranford et al, 2019; Incio et al, 2016a; Incio et al, 2016b; Khasawneh et al, 2009; Qureshi et al, 2020; Tiwari et al, 2019; Wang et al, 2019). However, the immune cell compartment of the tumor microenvironment and its possible impact on obesity-induced tumors has not been systematically characterized at the single-cell level.

Cancer progression is an evolutionary process where the fitness of cancer cells is dependent on reciprocal interactions between tumor -intrinsic and -extrinsic factors, including immune cells. In particular, CD8 T cells are central to tumor immunity and tumor-infiltrating CD8 T cells are associated with increased patient survival (Liu et al, 2012; Mahmoud et al, 2012; Martínez-Lostao et al, 2015). In the tumor microenvironment, MDSC possess strong T cell suppressive capacity inhibiting T cell function and proliferation (Bronte et al, 2016; Parker et al, 2015). Tumor associated macrophages with the often oversimplified M1/M2 characterization, along with T cells, NK cells, DCs, B cells, and eosinophils have complex and often inconsistent functions in cancer (Gonzalez et al, 2018; Noy & Pollard, 2014; Varricchi et al, 2017; Wylie et al, 2019).

Because of this intricate immune cell composition, conventional methods fail to reach the number of parameters required to profile the tumor immune microenvironment. High dimensional single cell approaches, such as mass cytometry, enable the simultaneous characterization of these varied cell types with multi-dimensional resolution.

Suspension mass cytometry (CyTOF) analysis of dissociated tumors can detect the multiple immune cell subsets required for an in-depth tumor immunotyping, but these data sets tend to be large and time consuming to collect. It is common to have data sets in multiple batches that are prepared, stained, and collected on different days due to the length of time needed to process and stain the samples and subsequently collect the data. Palladium-based metal barcoding of up to 20 samples has allowed for simultaneous collection of multiple samples without run/batch differences (Zunder et al, 2015). This is a great improvement; however many studies are composed of more than 20 samples. Therefore, it is common to end up with data in multiple batches that need to be compared. Having a common, or anchor, sample used in every batch can assist in the removal of batch effects but is not always available.

Here we have implemented a robust mass cytometry analysis pipeline to correct for batch effects between unanchored batches with multistep clustering to maximize phenotyping and minimize bias. We have immunophenotyped tumor immune infiltrate from two syngeneic pancreatic and three syngeneic breast cancer models and present that data as an immunotype atlas containing 21 immune cell metaclusters present across the five tumor models. Additionally, we report immunotyping of tumors grown in obese mice for all five models. Our findings demonstrate that the tumor immune infiltrate composition is highly model, and cancer type specific. One model, E0771 tumors, had significant immune cell differences between lean and obese mice. This breast cancer model showed an increase in G-MDSC and PD-L1+ DCs and a decrease in CD8 T cells in tumors from obese mice, making it a clinically relevant model (Jin & Hu, 2020; Mahmoud et al, 2012; Mahmoud et al, 2011).

## Results

### Tumor immune infiltrating cells were identified for 7 CyTOF batches from obese and non-obese mice

To mimic an obese environment, both male and female C57Bl/6 wild type mice were fed HFD or chow for 10 weeks prior to orthotopic transplantation of cancer cells (Fig 1A). High fat feeding resulted in higher body weights compared to the chow fed mice for male and female mice (Fig 1B). Murine cancer cells were then implanted orthotopically and tumors allowed to form while the mice were kept on their respective diets. To compile a model-independent systematic analysis of the tumor immune effects of the obese environment we investigated five syngeneic cell line tumor models in 2 cancer types (mammary adenocarcinoma: E0771, TeLi, and Wnt1; pancreatic ductal adenocarcinoma: C11 and UN-KC; Table 1). Consistently across the 7 batches, tumors grown in the obese environment displayed larger tumor mass than those grown in non-obese environments (Fig 1C).

**Figure 1.**
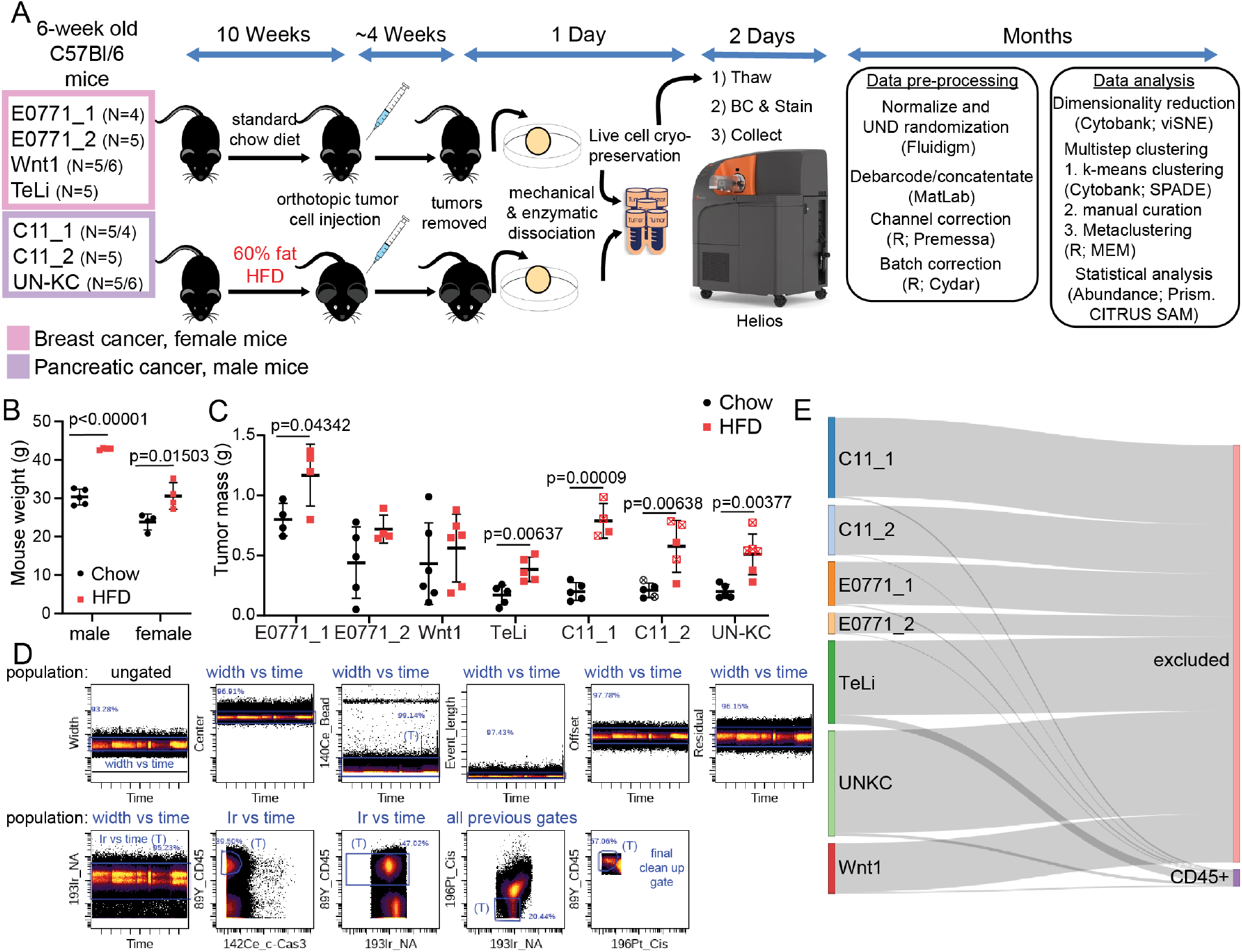
Experimental design and analysis pipeline for mass cytometry data for immune infiltrate of 7 CyTOF batches from 5 murine tumor models. A Cartoon and timeline of experimental design, data collection, data preprocessing, and analysis. B Representative mouse weights for male and female C57Bl/6 mice on chow and HFD. Weights were collected at age 16 weeks before tumor cell injection. C Tumor masses from the 7 batches for all experimental tumors run on CyTOF. D Representative gating strategy for identifying live CD45+ tumor infiltrating leukocytes. E Sankey plot visualization of CD45+ live cells out of total raw events collected for each experimental batch.

**Table 1.**
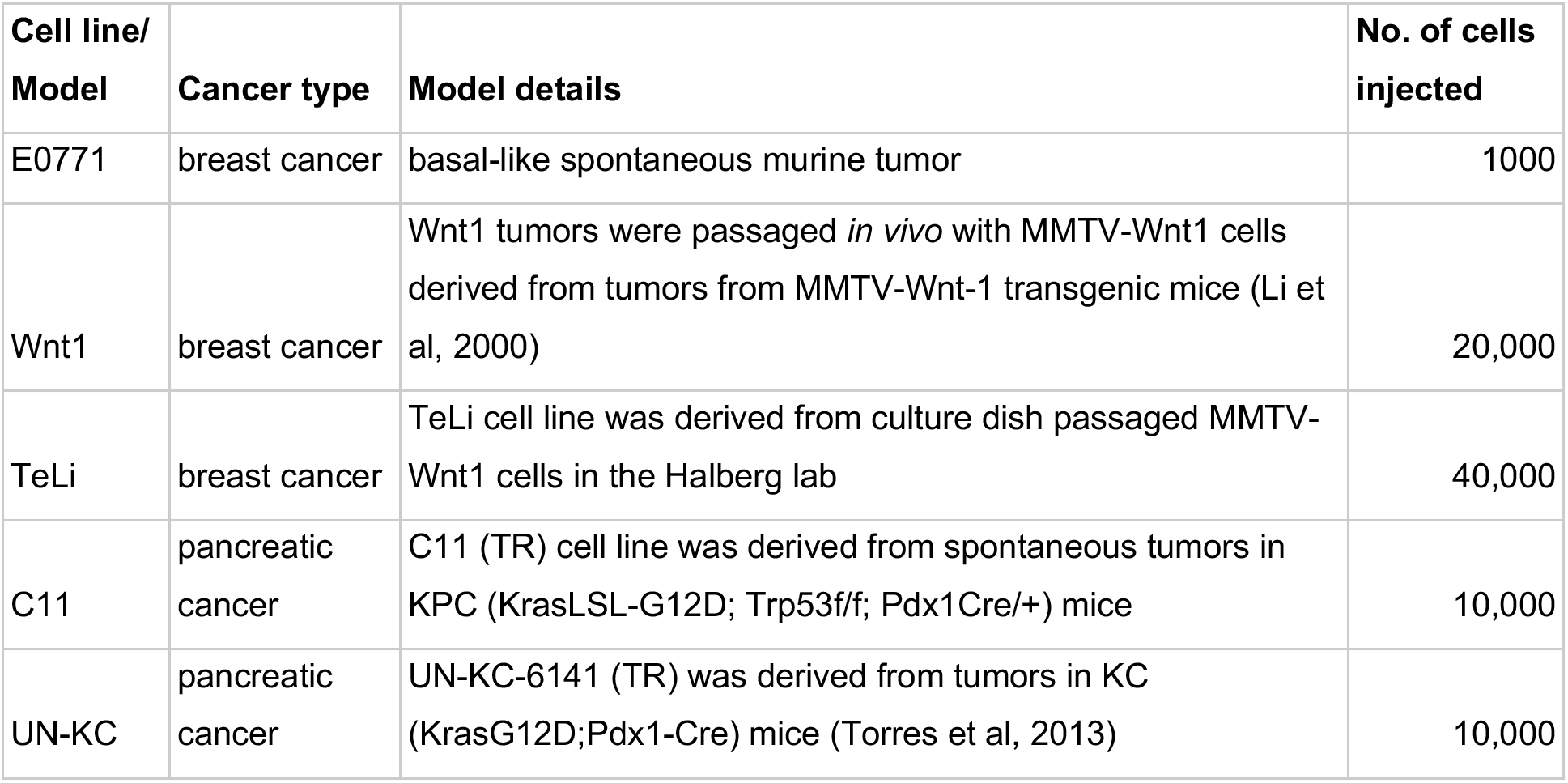
Cancer models. Tumor cells are syngeneic with C57Bl/6 mice and were orthotopically implanted into immunocompetent C57Bl/6 mice. E0771 cells and C11 cells were additionally implanted into TKO mice. Breast cancer cells were injected into female mice and pancreatic cancer cells were injected into male mice. TR: triple reporter-labeled

Tumors grown in obese or non-obese environments were then dissociated and immunophenotyped with a 36-marker immune-focused panel (Table 2) and analyzed by Helios CyTOF mass cytometry (Fig 1A). Single cell data from the five tumor models was collected in 7 individually barcoded batches (Fig 1A) allowing for samples in a single batch to be readily compared. FCS files were pre-processed and live CD45+ cells were gated separately for each batch (Fig 1A, D). 2.6 to 13.2 million total raw events were collected for each experimental cohort (see Table 3 for details). Tumor samples were not enriched for immune cells in advance to avoid experimental bias and preliminary data indicated that the immune cells comprise roughly 5% of the total collected events. With this in mind, the goal was to collect 100,000 to 1 million events per barcoded sample. Because of the lower percentage of target cells, the maximum number of barcodes used for a single batch was limited to 14. The percent immune cells per batch ended up ranging from approximately 1% to 12% (Table 3). In all tumor models, except TeLi, less than 5% of events collected were gated as live CD45+ single cells (Fig 1E).

**Table 2:** Panel of antibodies for mass cytometry staining All compounds were purchased from Fluidigm *IC = Intracellular (v) Markers used as parameters when running viSNE and CITRUS (m) Markers used as parameters when running MEM IRF4-155Gd (IC) was not included in the Wnt1 batch and was therefore removed from analysis. The staining intensity was low in the other stained batches.

| **Isotope tag** | **Target** | **Notes/expression/lineage** | **Target site** | **Dilution factor** | **Clone/ (Cat. Nr.)** |
| --- | --- | --- | --- | --- | --- |
| 089Y | CD45 | Pan leukocyte marker | Surface | 100 | 30F11 |
| 141Pr | Ly-6G (v,m) | Myeloid lineage, G-MDSC | Surface | 400 | 1A8 |
| 142Nd | c-Caspase 3 | Apoptosis marker | IC* | 500 | D3E9 |
| 143Nd | CD357/GITR (v,m) | Activation marker for T cells | Surface | 100 | DTA1 |
| 144Nd | MHC I/H-2Db | MHC-I on C57BL/6 | Surface | 600 | 28-14-8 |
| 145Nd | CD4 (v,m) | T helper cells | Surface | 100 | RM45 |
| 146Nd | F4/80 (v,m) | Macrophage marker | Surface | 200 | BM8 |
| 147Sm | CD36/FAT (v,m) | Scavenger receptor | Surface | 200 | No. 72-1 |
| 148Nd | CD11b/Mac-1 (v,m) | Macrophages and MDSC | Surface | 500 | M1/70 |
| 149Sm | CD19 (v,m) | B cells | Surface | 100 | 6D5 |
| 150Nd | Ly-6C (v,m) | Ly-6C+ monocytes, M-MDSC | Surface | 200 | HK1.4 |
| 151Eu | CD25/IL-2R (v) | T regs | Surface | 100 | 3C7 |
| 152Sm | CD3e (v,m) | T cells | Surface | 100 | 1452C11 |
| 153Eu | CD274/PD-L1 (m) | Ligand of PD-1 (marker of activation or exhaustion) | Surface | 400 | 10F.9G2 |
| 154Sm | CD73 (v,m) | B, T cells and macrophages. | Surface | 700 | TY/11.8 |
| 156Gd | CD14 (v,m) | LPS co-receptor on monocytes and macrophages | Surface | 100 | Sa142 |
| 158Gd | CD9 (v) | Cancer cells, eosinophils, activated T cells | Surface | 1000 | KMC8 |
| 159Tb | CD279/PD-1 (m) | Immune-inhibitory receptor on T cells, activation or exhaustion | Surface | 200 | J43 |
| 160Gd | CD5 | T cells and some B cells | Surface | 500 | 537.3 |
| 161Dy | CD40 (v,m) | Mature B cells, DCs and M1-like macrophages | Surface | 200 | HM403 |
| 162Dy | CD11c (v,m) | Dendritic cells, NK cells and macrophages | Surface | 300 | N418 |
| 163Dy | CD54/ICAM-1 (m) | B cell, T cell, monocytes, macrophages, NK and DCs | Surface | 600 | YN1/1.7.4 |
| 164Dy | Ly-6A/E/Sca-1 (v,m) | Hematopoietic stem cells and activated T cells | Surface | 900 | D7 |
| 165Ho | CD161/NK1.1 (v,m) | NK cells and NKT cells | Surface | 100 | PK136 |
| 166Er | CD326/EpCAM (v) | Cell adhesion in inflammation and carcinogenesis | Surface | 300 | G8.8 |
| 167Er | CD335/Nkp46 (v) | NK cells | Surface | 100 | 29A1.4 |
| 168Er | CD8a (v,m) | Cytotoxic T cells | Surface | 200 | 53-6.7 |
| 169Tm | CD206/MMR (v,m) | Mannose receptor, M2-like macrophages | IC/Surface | 200 | C068C2 |
| 170Er | CD169/Siglec-1 (v,m) | Macrophage restricted | Surface | 300 | 3D6.112 |
| 171Yb | CD44 (v,m) | Cell-cell interactions, adhesion, homing and migration | Surface | 1000 | IM7 |
| 172Yb | CD86 (v,m) | T cell responses binds CTLA-4 and CD28, M1-like macrophages | Surface | 200 | GL1 |
| 173Yb | CD117/c-kit | Mast cells and hematopoietic stem cells | Surface | 100 | 2B8 |
| 174Yb | CD223/LAG-3 | T cell activation and NK cells | Surface | 100 | C9B7W |

Table 2: Panel of antibodies for mass cytometry staining
| Isotope tag | Target | Notes/expression/lineage | Target site | Dilution factor | Clone/ (Cat. Nr.) |
| --- | --- | --- | --- | --- | --- |
| 175Lu | CD127/IL-7Ra (v,m) | Memory T cells | Surface | 100 | A7R34 |
| 176Yb | CD278/ICOS (m) | Activated T cells | Surface | 100 | 7E.17G9 |
| 209Bi | MHC II/I-A/I-E (v,m) | APCs | Surface | 1000 | M5/114.15.2 |
| Ir191/193 | Nucleic acid | Iridium DNA intercalator | IC |  | (201192B) |
| Pt196_Cis | Membrane exclusion | Cisplatin, cell viability stain | IC |  | (201064) |
| Palladium | Non-specific covalent | BarCode kit, 102, 104:106, 108, 110 | IC |  | (201060) |

**Table 3.** CyTOF batches. All batches were collected on the Helios CyTOF machine in the flow cytometry core at the University of Bergen. 69 channels were collected for all batches including the Helios Gaussian channels. Values in () indicate the tumors with more than 5000 immune cells that were used for downstream analysis when not all tumors could be analyzed.

| Batch | days of tumor growth | non-obese tumors analyzed by CyTOF | Obese tumors analyzed by CyTOF | total barcoded samples including controls | Total raw FCS files collected | Raw data size in GB | total raw events | total CD45+ cells including controls and low count samples | Percent of CD45/total raw events |
| --- | --- | --- | --- | --- | --- | --- | --- | --- | --- |
| E0771_1 | 31 | 4 | 4 | 9 | 7 | 4.27 | 5479930 | 180207 | 3.3 |
| E0771_2 | 23 | 5 | 4 | 10 | 9 | 2.05 | 2636328 | 100935 | 3.8 |
| TeLi | 41 | 5 | 5 | 11 | 8 | 7.43 | 9544773 | 1102767 | 11.6 |
| Wnt1 | 27 | 6 | 6 | 14 | 17 | 4.89 | 6284192 | 193650 | 3.1 |
| C11_1 | 31 | 5 | 4(1) | 10 | 10 | 7.93 | 10180942 | 207180 | 2.0 |
| C11_2 | 32 | 5(3) | 5(2) | 11 | 5 | 4.92 | 6319718 | 54317 | 0.9 |
| UN-KC | 26 | 5 | 6(2) | 14 | 11 | 10.3 | 13193861 | 369684 | 2.8 |

### Unanchored range-based batch correction enabled successful co-analysis of CyTOF data from multiple barcoded batches

Streamlined analysis between batches is limited by technical issues such as staining intensity and machine variation (Kleinsteuber et al, 2016; Leipold, 2015; Leipold et al, 2018; Schuyler et al, 2019). To enable streamlined cross-analysis between tumor batches and models we tested 3 unanchored batch correction algorithms available through Cydar (Lun ATL, 2017). To test the robustness of the batch correction, we combined our 7 batches with 2 additional batches of tumor immune cells from 4T1 syngeneic tumors grown in BalbC mice with a different cell history and staining panel. We first tested batch correction using warp, quantile, and range batch correction approaches on the 9 testing batches with 18 shared markers (Fig 1A, EV1, Table 4). The need for batch correction can be seen in the variable distribution of CD11b and F4/80 positivity in the uncorrected plots (Fig 2A, EV1A). Pre batch correction, the CD11b signal is low for C11_1, Wnt1, and TeLi and high for both 4T1 batches. The signal intensity becomes more normalized with warp and range batch correction applied. Quantile batch correction, on the other hand, performed poorly and caused an increase in noise and a distortion of the density distribution in the samples, as indicated by the black arrows (Fig EV1A). This distortion of the data is most clearly observed for Ly6-G, where the need for batch correction was minimal as seen by the closely aligned peaks in the uncorrected Ly6-G plot. Warp and range correction had similar performance to each other and were therefore further evaluated with the 9 testing batches (Fig EV1B-D, 2A).

**Table 4.**
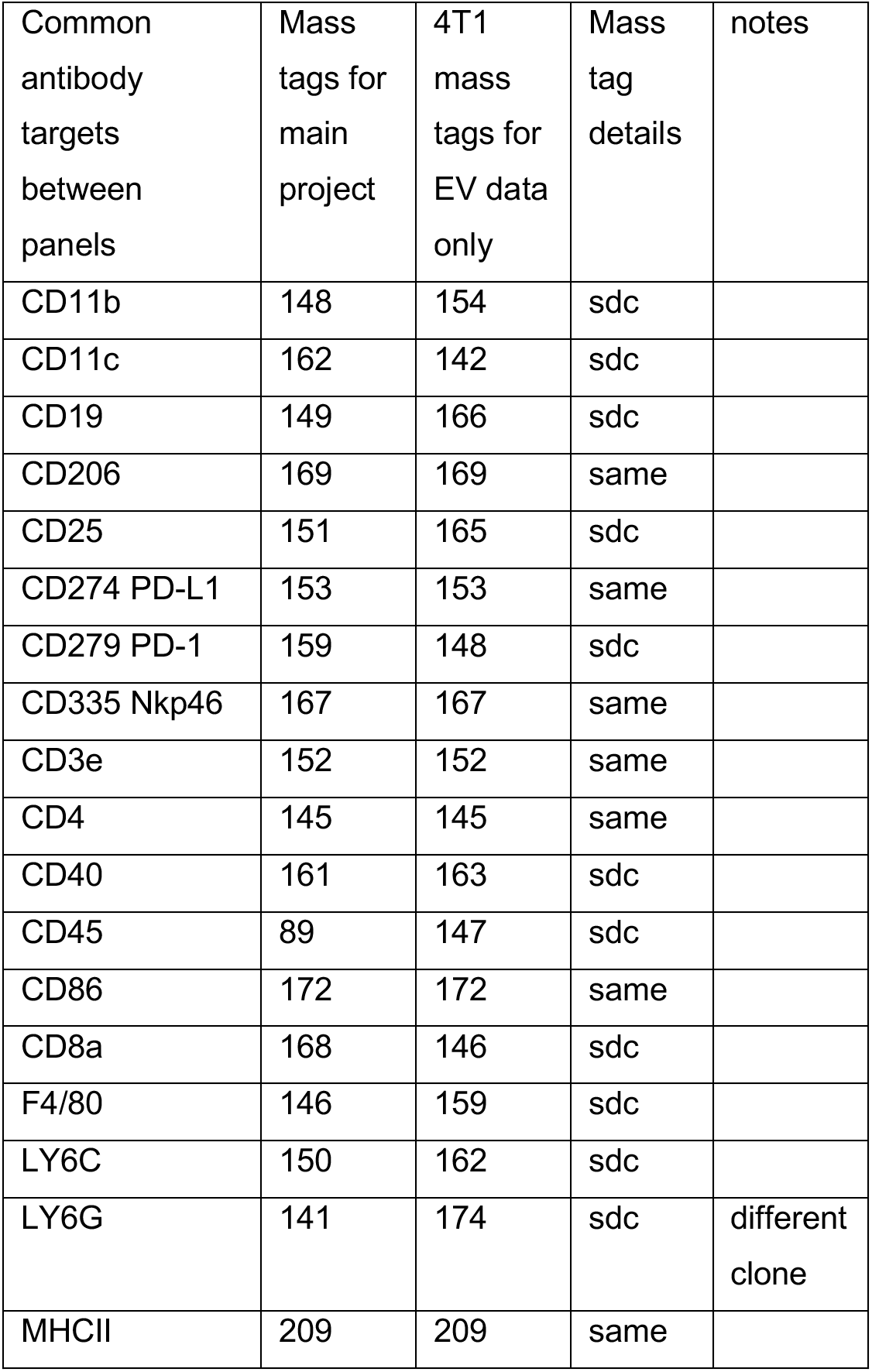
Common channels used for batch correction testing with 9 batches sdc – shared antibody, different channel

| Common antibody targets between panels | Mass tags for main project | 4T1 mass tags for EV data only | Mass tag details | notes |
| --- | --- | --- | --- | --- |
| CD11b | 148 | 154 | sdc |  |
| CD11c | 162 | 142 | sdc |  |
| CD19 | 149 | 166 | sdc |  |
| CD206 | 169 | 169 | same |  |
| CD25 | 151 | 165 | sdc |  |
| CD274 PD-L1 | 153 | 153 | same |  |
| CD279 PD-1 | 159 | 148 | sdc |  |
| CD335 Nkp46 | 167 | 167 | same |  |
| CD3e | 152 | 152 | same |  |
| CD4 | 145 | 145 | same |  |
| CD40 | 161 | 163 | sdc |  |
| CD45 | 89 | 147 | sdc |  |
| CD86 | 172 | 172 | same |  |
| CD8a | 168 | 146 | sdc |  |
| F4/80 | 146 | 159 | sdc |  |
| LY6C | 150 | 162 | sdc |  |
| LY6G | 141 | 174 | sdc | different clone |
| MHCII | 209 | 209 | same |  |

**Figure 2.**
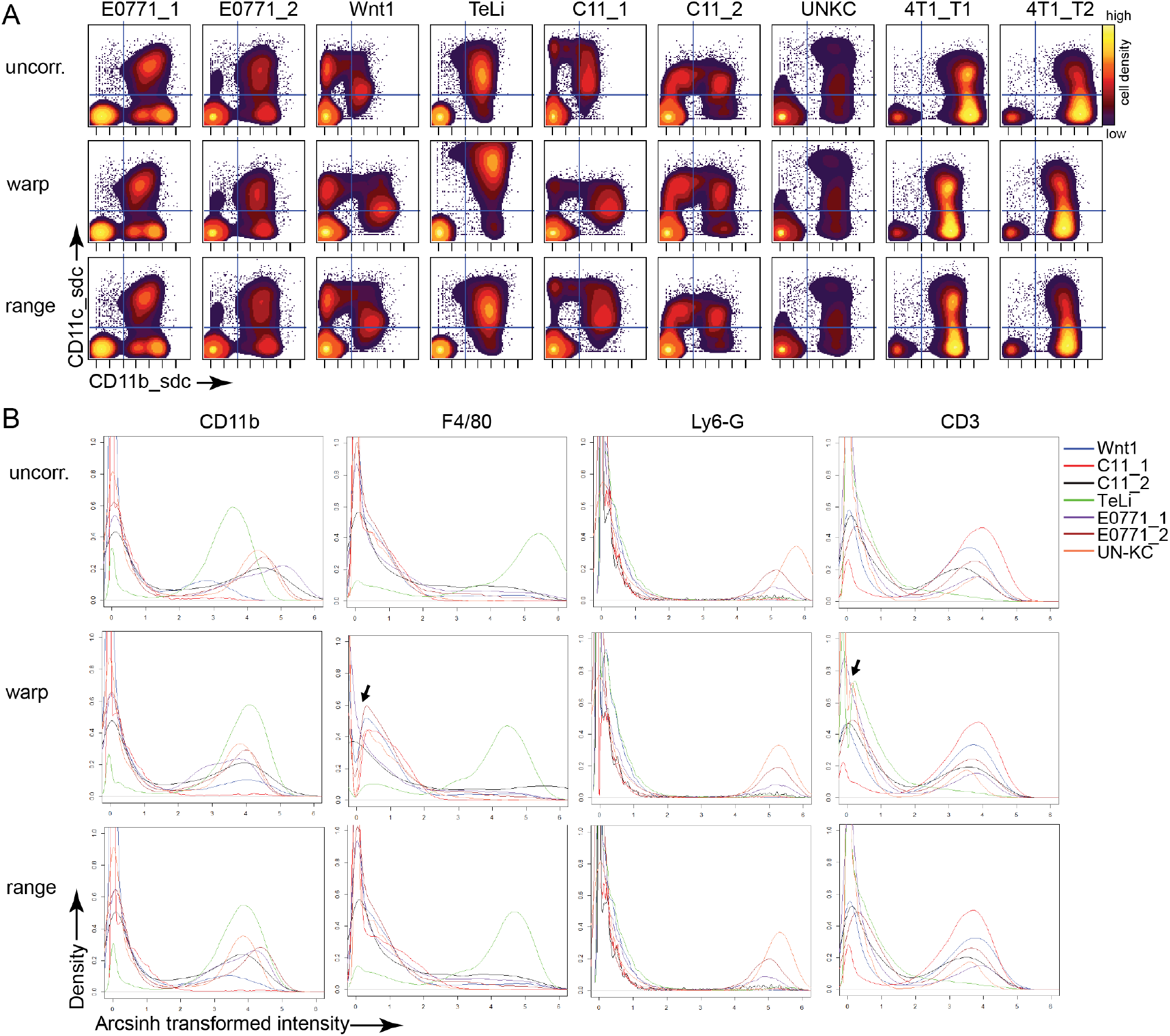
Batch correction algorithm testing. All plots were generated from normalized arcsinh transformed live CD45+ cells. Transformed datasets were warp and range corrected resulting in three datasets, incluing the uncorrected (uncorr.) dataset. A Biaxial contour density plots from the 9 testing batches with 18 common markers. The files displayed are the first chow/control sample from each batch. The quadrant gate is shown to assist in visual comparison between plots. B Cydar batch correction density plots of 4 representative markers showing the 3rd file from each of the 7 experimental batches with 36 common markers total. Black arrows indicate a gap in the density near zero created by the warp correction algorithm.

Warp and range corrected datasets resulted in almost identical viSNE maps when run together (Fig EV1B) and separately with the same seed (Fig EV1C). The marker signal intensity varied somewhat between warp and range corrected files run in the same viSNE but the cell placement on the viSNE map was almost identical (Fig EV1B). This alignment of cell placement can also be seen in the linear regression of the tSNE1 and tSNE2 channels between warp and range (Fig EV1D). While the warp correction algorithm closely aligns the density peaks (Fig EV1A), it also added distortion artifacts to the data. The warp correction distorted the CD11c signal in the TeLi plot making the signal higher than in the uncorrected data and higher than in the other batches (Fig 2A, EV1C black arrow). The range corrected plots showed a more consistent max intensity for CD11c. The warp correction also resulted in an unexplained bunching artifact for the E0771_2 plot wherein the cells in the lower left of the map were stacked in a small area (Fig EV1C pink arrow). To make the final determination between warp and range correction, the 4T1 batches were removed and the 7 experimental batches were batch corrected by warp and range methods for 35 markers and the density plots for key markers were evaluated (Fig 2B). For several markers, including F4/80 and CD3, the warp correction created an artificial gap in the data around zero as indicated by the black arrows (Fig 2B). Because of this and previously mentioned warping artifacts, Cydar’s range correction algorithm was chosen as the batch correction method for the 7 experimental batches. With successful batch correction, the data from the 5 models could then be analyzed in concert and a uniform analysis pipeline was implemented.

### viSNE-based immunotyping revealed diverse myeloid and lymphoid immune infiltrate across the tumor models

Having successfully batch corrected the experimental data, we next moved into the analysis pipeline beginning with dimensionality reduction using the Cytobank viSNE implementation of BH-tSNE (Amir el et al, 2013; Kotecha et al, 2010). 5206 live CD45+ immune cells from each of 57 experimental files and 10 control files from the 7 batches were run together in a single viSNE analysis with a final KL divergence of 4.75. 26 phenotyping markers were used to generate the viSNE map (see Table 2 for details). Cell density was plotted onto the viSNE map to visualize the overall cell distribution and heterogeneity of the tumor immune infiltrate (Fig 3A, B). The presence of the multiple density “islands” indicates a successful viSNE run and a diverse range of immune cells present across tumor types and diet groups. When assessing density differences between the plots, the largest differences appear to be between tumor models rather than between diet groups (Fig 3A).

**Figure 3.**
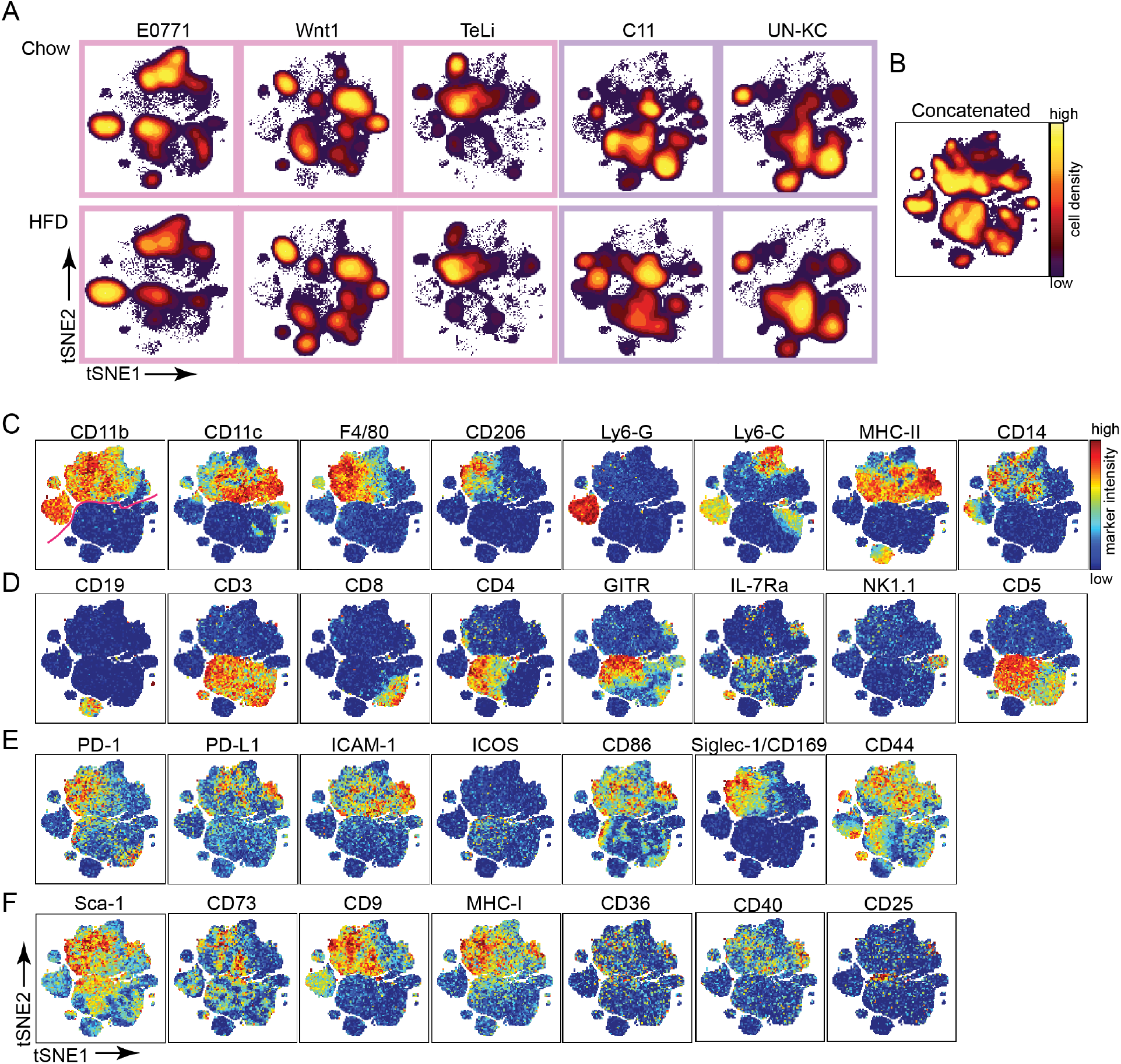
Immune infiltrate phenotyping using viSNE. All range corrected experimental files were run together in the same viSNE run to generate one universal viSNE map. A Cell density on viSNE plots for concatenated experimental files for chow and HFD groups in each cohort. B-F The total concatenated data was used to generate viSNE plots. B Density plot of total concatenated cells. C-F are viSNE plots showing marker intensity on a spectrum heat scale. Heat scales are specific to individual markers. C Marker heat for key myeloid phenotypic markers. The pink line on the CD11b plot indicates the phenotypic divide between myeloid and lymphoid cells. D Marker heat for key lymphocyte phenotypic markers. E Marker heat for activation/exhaustion markers. F Marker heat for additional phenotyping markers.

To enable visualization of the all the data in one plot, the files were concatenated (Fig 3B-F). Interrogating the viSNE map by marker intensity revealed the top half (denoted by pink line in first plot in Figure 3C) to be dominated by areas of distinct myeloid marker expression (CD11b, CD11c, F4/80, CD206, Ly6-G, Ly6-C, MHC-II, and CD14) indicating a diverse myeloid cell tumor infiltrate. Similarly, the bottom half of the map consists of diverse lymphocyte populations which can be identified by the marker heats for CD19, CD3, CD8, CD4, GITR, IL-7Ra, and NK1.1 (Fig 3D). Cells can be further characterized by looking at the marker expression for activation and exhaustion markers (Fig 3E) and additional phenotyping markers (Fig 3F) on the viSNE map.

### Cross model immunophenotyping of tumor immune cell metaclusters

We next wanted to identify and characterize the different immune cell subsets to compare cell type abundances between tumor models. To perform this analysis with minimal bias and without manual gating, we performed a series of clustering and curating steps to establish 21 biologically meaningful metaclusters. tSNE1 and tSNE2 were used as the input parameters for SPADE-based K-means clustering so that the results of the dimensionality reduction would be preserved in the clusters. We used a k of 40 for this first step to capture the immune diversity while reducing the possibility of under-clustering. The 40 clusters were then manually curated to reduce obvious over-clustering by combining clusters within small islands for G-MDSC and NK cells, resulting in a total of 37 clusters. To phenotypically characterize the 37 clusters and to hierarchically cluster them into metaclusters, we next used 26 markers to generate the MEM enrichment scores and to perform hierarchical clustering of the SPADE clusters and markers based on their MEM scores (Fig 4A, Table 2) (Diggins, 2016). MEM provides insight into how a cluster is positively (yellow) or negatively (blue) enriched for a specific marker compared to the other cells from the other clusters (Diggins et al, 2017). The cluster dendrogram (left hand side of Figure 4A) was used to create 21 metaclusters (MC) (Fig 4B). The MEM heat map and scores, median heat map (Fig EV2A), and viSNE plots (Fig 3B-E) were subsequently used to identify and label the 21 metaclusters. Automated clustering instead of manual gating within the CD45+ cells means that no cells were excluded or double counted and user bias was minimized. The multistep clustering pipeline (k-means clustering, followed by manual curation, and then dendrogram metaclustering) ensured accurate placement for each cell while minimizing over- and under-clustering. CD11b, F4/80, MHC-II, CD11c, Ly6-C, CD3, and GITR had the largest contribution to the MEM hierarchical clustering as determined by the marker dendrogram at the top of Figure 4A. These are bright markers present on many cells with large expression differences between the clusters.

**Figure 4.**
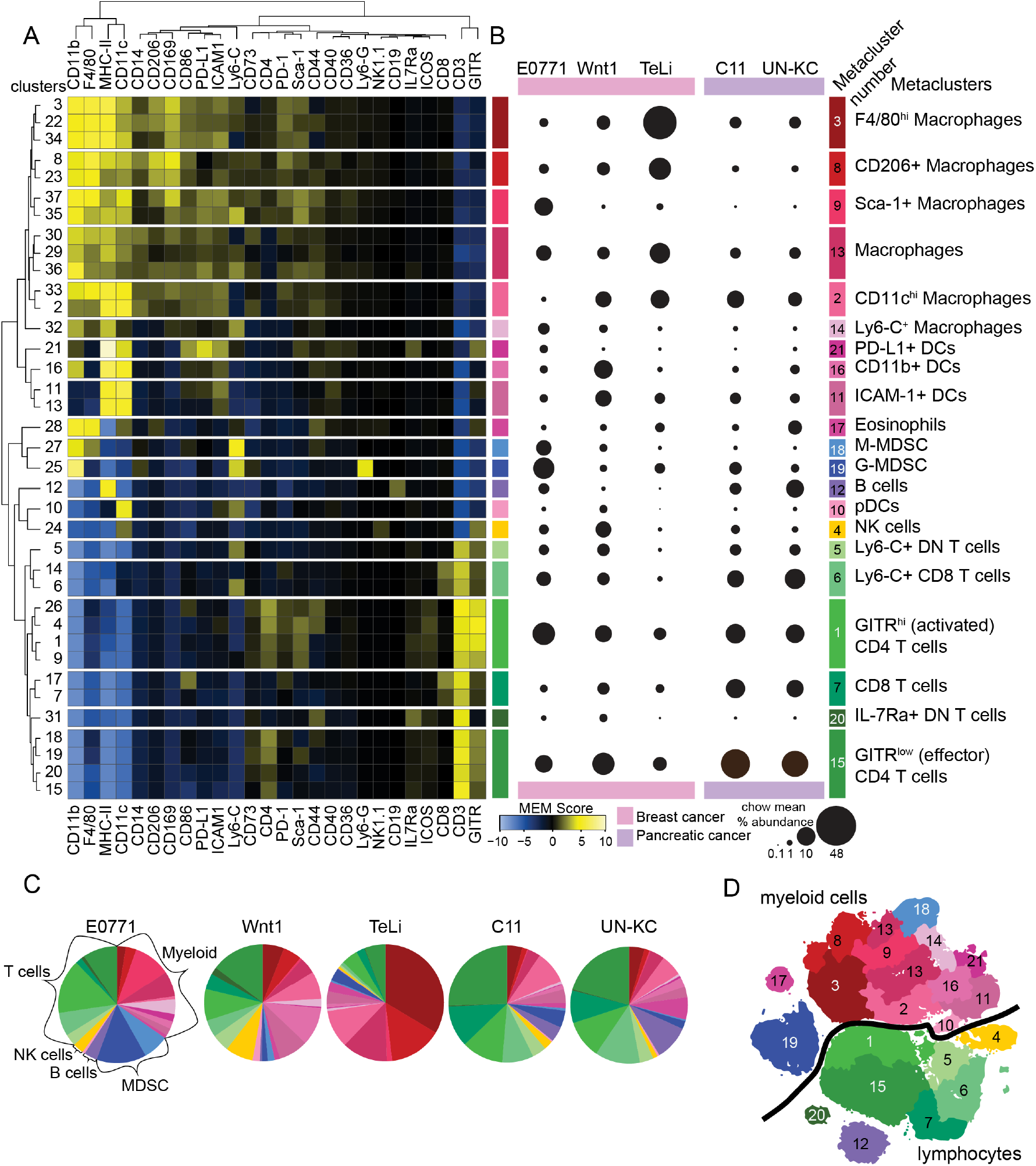
Immune infiltrate metacluster characterization of murine breast and pancreatic cancers. A Hierarchical clustering of MEM scores for 37 curated clusters and 26 markers. The dendrogram on the left was used to create 21 metaclusters. MEM scores and marker heat were used to label the metaclusters (labels in B). B Bubble graph using area to show the mean percent abundance out of the total CD45+ cells for each metacluster for immune infiltrate of chow-fed non-obese each tumor type. C Pie charts showing mean percent abundance of the 21 metaclusters across the 5 models for the chow tumors. D Annotated viSNE map of concatenated data showing the metaclusters by number. The black line indicates the divide between myeloid and lymphoid lineage cells/clusters.

This analysis pipeline allowed us to confidently identify 6 macrophage, 6 T-cell, 4 DC, and 2 MDSC metaclusters in addition to B cells, NK cells, and eosinophils. All samples were represented in all 21 of the metaclusters. An overview of the mean percent metacluster abundance for the non-obese group in each model is shown in the bubble graph (Fig 4B) and immune cell populations from the chow control groups in the different tumors are displayed as pie charts in Figure 4C. The concatenated metacluster data was plotted onto the viSNE map for visualization (Fig 4D).

Of the six macrophage metaclusters, MC8 was M2-like and MC2 was M1-like in phenotype. The CD206+ macrophages in MC8 were also enriched for CD14 and CD169 and had decreased expression of MHC-II. The M1-like CD11c+ macrophages (MC2) were more abundant in pancreatic cancer models and in the Wnt1 breast cancer model. The M2-like macrophages (MC8) were more abundant in the E0771 and TeLi models (Fig 4B). The other macrophage subsets are discussed below. The E0771 tumors had the largest abundance of Sca-1+ macrophages (MC9) with approximately 10% while they were minimally observed in the other models. MC3 macrophages were especially enriched for F4/80 while MC13 macrophages had no strong identifying markers. TeLi tumor infiltrate was dominated by macrophages, specifically from MC3 F4/80^hi^ macrophages, with 73% of TeLi chow immune infiltrate falling into the 6-macrophage metaclusters (Fig 4B, C). The other tumor models had a more diverse immune infiltrate makeup (Fig 4B, C). MC14 was a small subset of Ly6-C+ macrophages with the greatest abundance in the E0771 and Wnt1 models. The breast cancer tumor immune infiltrate was dominated by myeloid cells while pancreatic cancer was dominated by T cells (Fig EV2B, C).

Of the six T cell metaclusters, two were CD8+, two were CD4+, and two were double negative (DN). The CD8+ T cell metaclusters MC6 and MC7 were touching on the viSNE map and quite similar in phenotype (Fig 4A, B, D). MC6 contained Ly6-C+ and Sca-1+ CD8 T cells. MC6 and MC7 were most abundant in the pancreatic tumor models, with C11 average chow infiltrate at 9% and 11% and UNKC chow infiltrate at 12% and 9%, respectively (Fig 4). Both CD8 T cell subsets were present across tumor models and expressed similar levels of PD-1 (Fig 3E). The CD4 T cells fell into an activated GITR^hi^ subset (MC1) and an effector subset that was GITR^low^ (MC15). The DN T cell metaclusters were phenotypically distinct on the viSNE map and in the MEM heatmap with MC5 consisting of L6-C+ DN T cells and MC20 consisting of IL-7Ra+ DN T cells (Fig 4A, B, D).

Four Dendritic cell (DC) metaclusters were identified in the five tumor models. DCs in the four metaclusters were CD11c+, F4/80-, CD206-, and CD14-. MC21 was composed PD-L1+ DCs that were also enriched in MHC-II, CD86, ICAM-1, IL7Ra, and GITR (Fig 4A, EV2A) (ten Broeke et al, 2013). MC21 was a small metacluster with less than 1% abundance in most cohorts and the largest abundance in E0771, which was less than 4% (Fig 4B). Plasmacytoid DCs (pDCs) (MC10) were the least abundant of the DC metaclusters across all models, less than 2% for Wnt1 tumors and less than 1% for all other models (Fig 4B, C). An ICAM-1+ DC metacluster (MC11) was the second largest DC metacluster for the Wnt1 model and the largest metacluster for the other four models. Wnt1 had the most DCs of the tumor models with DCs primarily falling into MC16 and MC11. MC16 was composed of CD11b+ DCs and was the most abundant metacluster for the Wnt1 model with approximately 8% for chow tumors.

MDSC were subsetted into two metaclusters, G-MDSC (also known as PMN-MDSC) in MC19 and M-MDSC in MC18. Both cell types are CD11b+; G-MDSC are Ly6-G+ and Ly6-C^low^ while M-MDSC are Ly6-C+ and Ly6-G− (Bronte et al, 2016). MDSC are known suppressive cells with G-MDSCs resembling granulocytes and M-MDSCs resembling monocytes. The G-MDSCs in MC19 were particularly abundant in the E0771 chow tumors (Fig 4B, C).

For the non-obese groups, this unbiased analysis highlights a remarkable heterogeneity between the syngeneic models. The overall myeloid cell abundance was 50% or greater for the breast cancer models and less than 40% for the pancreatic cancer models. In the pancreatic tumor models, over 50% of the infiltrate was T cells. (Fig 4C, EV2C). A comparison of breast and pancreatic cancer immune infiltrate for the non-obese groups revealed multiple significant differences in metacluster abundance (Fig EV2B). However, a limitation is that additional variables such as sex, cancer cell line, and tumor location have not been controlled for in this analysis.

### The obese microenvironment leads to model specific alterations in immune cell populations

Having defined 21 phenotypically relevant metaclusters, we next asked if an obese environment was associated with differential abundance in immune cell types. Metacluster percentages out of total tumor infiltrating immune cells were plotted for chow and HFD tumors for each model (Fig 5A, B).

**Figure 5.**
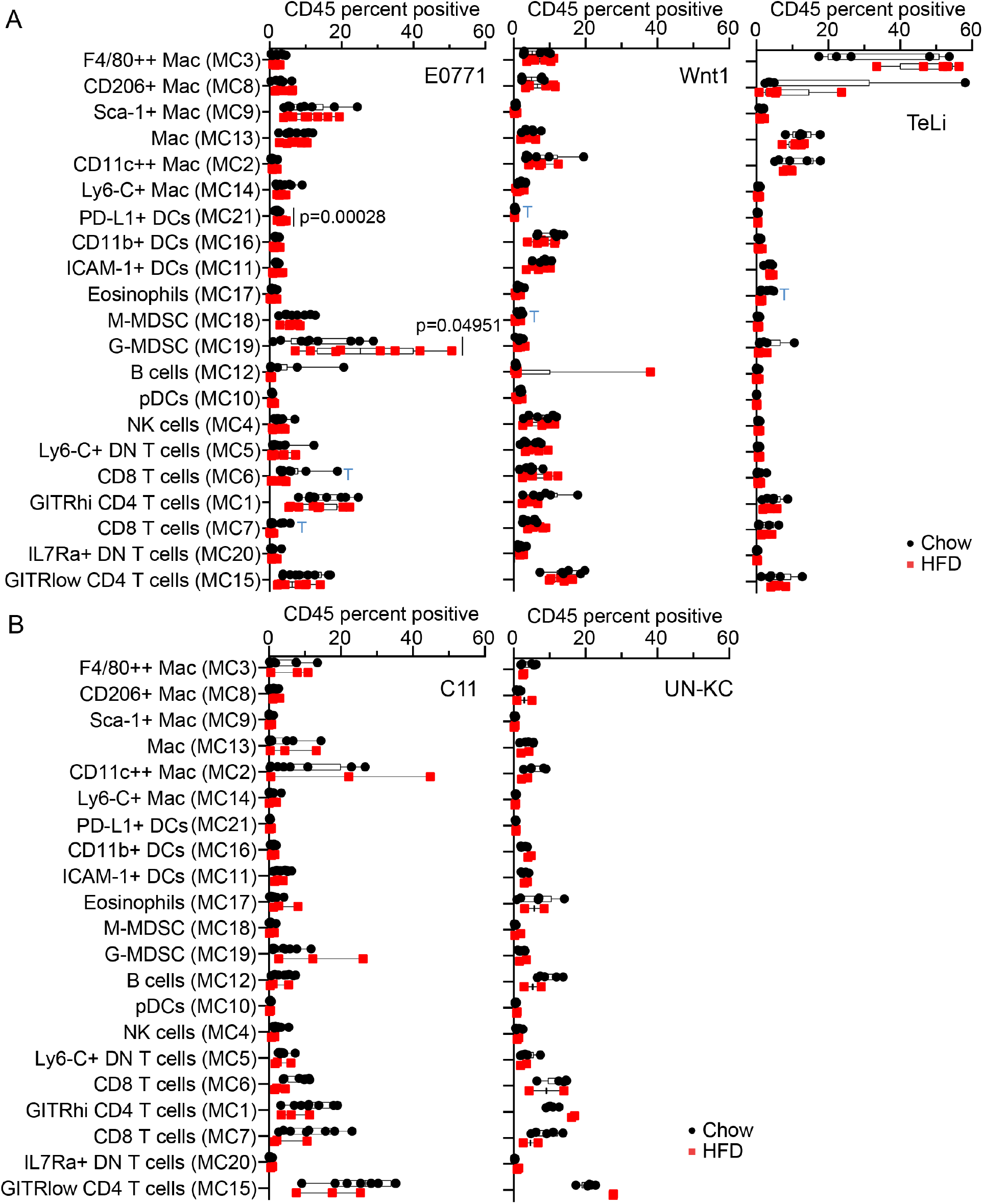
Analysis and quantification of metacluster abundance differences between chow and HFD. A Min to max box and whiskers plots with all points showing comparing chow and HFD metaclusters for the breast cancer cohorts. Significant p values are shown in the figure. The blue T indicates p values less than 0.07 that are trending towards significance. B Box and whiskers plots for pancreatic cancer cohorts. There were too few HFD tumors with live immune cells so statistics could not be performed for those cohorts.

Surprisingly, the two Wnt1-driven mammary tumor models (Wnt1 and TeLi) displayed no significant differences between chow and HFD in tumor immune infiltrate (Fig 5A). However, in the E0771 breast cancer model, we observed significant differences in a small PD-L1+ DC metacluster (MC21) and the large G-MDSC metacluster (MC19), both showing increased abundance in HFD (Fig 5A). The E0771 tumors from obese mice contained the highest percentage of G-MDSC which was significantly more than the tumors from non-obese mice (Fig 5A). Both CD8 T cell metaclusters (MC6 and MC7) were trending towards a decrease in HFD (Fig 5A). In the pancreatic cancer models, the obese tumor microenvironment did not lead to any major alterations in the tumor immune infiltrate.

### Diet-induced obesity led to an increased abundance of G-MDSC and a decrease in CD8 T cells in the E0771 triple negative breast cancer model

To further explore the connection between the obese microenvironment and immune infiltrate in the E0771 model, we examined the ratio between CD4 and CD8 T cells. This ratio has been used in peripheral blood and tumor tissue as a measure of immune health (Das et al, 2018). In breast cancer an elevated CD4/CD8 ratio has been associated with tumor progression and poor survival (Wang et al, 2017; Yang et al, 2017). Here we found that the CD4/CD8 ratio was higher in tumors that evolved in obese compared to non-obese mice in the E0771 model (Fig 6A). Further, the CD8 T cell percentage out of the total T cells was significantly decreased in HFD tumors in the E0771 model (Fig 6B). We did not detect differences for the CD4 T cell population in the E0771 model (Fig EV3A). For E0771, the total T cells out of the CD45+ cells were trending towards a decrease in HFD but the results were not significant (Fig EV3B).

**Figure 6.**
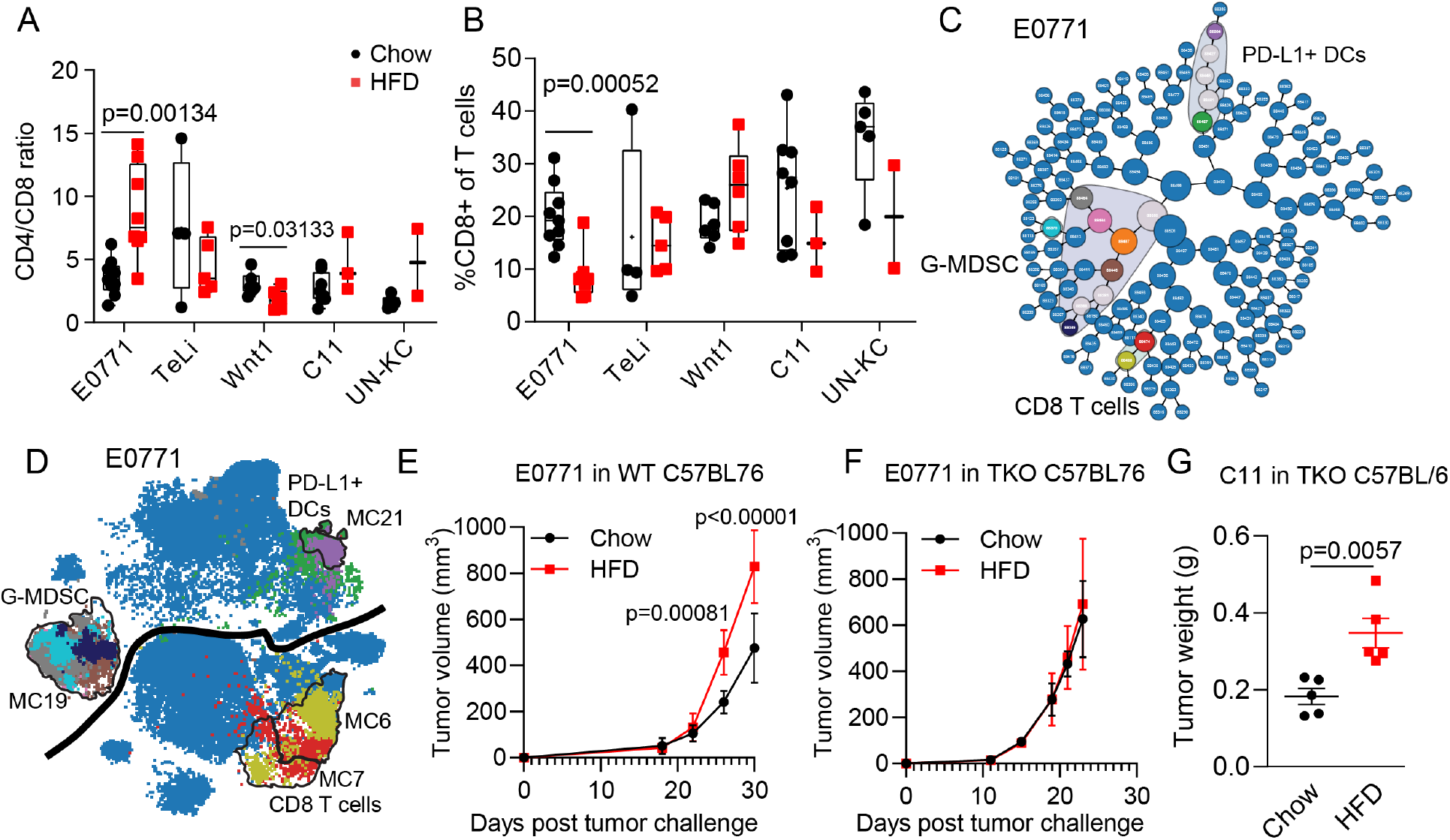
CD8 T cells were decreased in the HFD DIO E0771 model of TNBC and tumor growth advantage was lost when the T cell compartment was lost in the TKO model. A Box and whiskers plots of CD4/CD8 ratio for each tumor model. B Percent of CD8 T cells out of total T cells. C CITRUS SAM results for E0771 model. CITRUS Clusters that are significantly different between chow and HFD are not blue and are circled with a gray background. D Select significant CITRUS clusters were plotted back onto the viSNE map. Plotted clusters are color coded to match the CITRUS plot in C. viSNE data shown is concatenated data for the 10 CITRUS clusters and total cell numbers for the E0771 model. E-F Tumor growth volume over time for E0771 WT (E), and E0771 TKO (F) cohorts. G Final tumor masses for C11 tumors grown in TKO mice showing that the HFD tumor growth difference remains. The appropriate comparison is to the C11 tumor masses for C11_1 and C11_2 batches shown in Figure 1C.

To independently validate the E0771 findings, we next ran a CITRUS analysis on the E0771 cohort (Bruggner et al, 2014). Consistently, the CITRUS SAM model found significant cell abundance differences for the three groups described above, G-MDSC, CD8 T cells, and MHC-II^hi^ DCs (Fig 6C, D). A subset of the significant clusters were mapped onto the viSNE map for cell type identification and visualization (Fig 6D). The abundance differences were in the same direction as identified in the metaclusters in Figure 5A, with G-MDSC and MHC-II^hi^ DCs increasing in HFD and CD8 T cells decreasing in HFD (Fig EV3C). G-MDSC have been reported to inhibit CD8 T cell function and proliferation (Youn & Gabrilovich, 2010). We thus hypothesized that the increased G-MDSC population was inhibiting CD8 T cell effector function and proliferation resulting in larger tumors in the obese E0771 model.

To functionally test this, we implanted E0771 cancer cells into the mammary gland of C57Bl/6 mice deficient in T-, B- and NK-cells (*Rag2*^−/−^::*CD47*^−/−^::*Il2rg*^−/−^; TKO) and compared the tumor growth to tumors in wild type (WT) C57Bl/6 mice. Interestingly, the tumor growth advantage observed in the obese environment of wild type mice, was abrogated in the immune deficient environment in the TKO mice (Fig 6E, F). Consistent with our immune profiling, this suggests E0771 cancer cells that evolve in the obese environment have the ability to attract G-MDSC to overcome T-cell cytotoxicity. This renders tumor growth independent of T cell infiltration in the obese state. In the non-obese state, however, E0771 tumor growth is affected by cytotoxic T cells and tumor growth is therefore enhanced in the non-obese TKO mice. In addition to the E0771 model, we also performed the TKO experiment in the C11 pancreas tumor model. Our immunotyping of this tumor did not suggest any deregulated immune populations in HFD (Fig 5B). Consistently, the C11 tumor growth advantage in the obese environment was sustained in the TKO model (Fig 6G). Combined, this demonstrates that in the E0771 model, tumor evolution in obese environments are linked to a functional differential immune interaction and that such interaction is highly model dependent.

## Discussion

Here we compared the tumor infiltrating immune cell populations from 7 unanchored batch corrected Helios CyTOF runs collected from two different tumor types adapted to obese and non-obese environments. The batches were then combined to form 5 tumor models for in depth immunotype analysis.

While anchored batch correction is the new gold standard for batch correction methods, there are many datasets that do not have this luxury. Our unanchored batch correction implementation here enabled the implementation of a single comprehensive analysis pipeline and provides a path forward for the streamlined analysis of other unanchored multi-batch mass cytometry datasets. Being able to analyze datasets together provides a great advantage over analyzing them in parallel. Batch corrected datasets can be gated, clustered, visualized, and statistically analyzed in unison making for stronger conclusions. Application of unanchored batch correction could allow for the meaningful reanalysis of previously collected data sets that may have been set aside due to batch effects and lack of anchor or reference samples.

The implementation of automated clustering approaches over manual gating have introduced a shift in cell subset classification rendering clustering less reliant on the+/− classification system traditionally used for characterizing cell types (Misharin et al, 2013; Zaynagetdinov et al, 2013). Cluster subsetting herein was performed with dimensionality reduction and a multistep clustering approach that minimized bias due to a lack of manual gating. For example, CD11b+ DCs (seen in MC16) have typically been described as F4/80 positive or negative (Misharin et al, 2013), but because we used F4/80 to classify cells, F4/80 positive cells are separated from F4/80 negative cells. Here, F4/80+ cells were classified as macrophages, eosinophils, or M-MDSC and all of the DC subsets were F4/80−.

Macrophages and DCs form a complex family of myeloid cells with overlapping functions and phenotypes (Misharin et al, 2013). Their highly plastic nature and tissue specific phenotypes make identification difficult (Leopold Wager & Wormley, 2014). We chose to name the metaclusters with the most likely cell type based on standard marker classifications and by the marker that most distinguished them from the other metaclusters of that cell type. Additionally, we included a wide range of phenotyping data in the figures to act as a resource for others to identify cell types of interest regardless of the name used to define them. This study identified multiple tumor infiltrating macrophage and DC subsets.

Macrophages, identified by high expression of CD11b, F4/80, and MHC-II, fell into 6 metaclusters representing macrophages of differing phenotypes and function. CD206 expression is associated with an M2 or pro-tumor phenotype (Haque et al, 2019; Nawaz et al, 2017). CD11c^hi^ macrophages (MC2) were phenotypically similar to an M1, known to be an anti-tumor phenotype macrophage (Gautier et al, 2012; Noy & Pollard, 2014; Zhu et al, 2017). The CD11c^hi^ macrophages were also high for MHC-II, indicating an activated state (ten Broeke et al, 2013). MC14 was composed of Ly6-C+ macrophages, which have been described in the literature as playing a detrimental role in multiple disease states (Gibbons et al, 2011; Kimball et al, 2018). The Ly6-C+ macrophages may also represent newly infiltrating monocytes which may further differentiate into other macrophage phenotypes (Movahedi et al, 2010; Noy & Pollard, 2014; Rahman et al, 2017). Sca-1+ macrophages (MC9) were additionally enriched for ICAM-1 and PD-L1. ICAM-1 has been reported to have anti-tumor effects while PD-L1 is associated with T cell inhibition with pro-tumor effects (Han et al, 2020; Patsoukis et al, 2012; Yang et al, 2015) Sca-1 expression is associated with stemness and a self-renewing state (Walasek et al, 2013), making this macrophage phenotype particularly complex.. These six metaclusters reveal the complexity of tumor associated macrophages and quantify their contributions to the tumor microenvironment.

It is interesting to note that the Wnt1 and TeLi immunotypes are so different with TeLi infiltrate being dominated by macrophages. These models only differ by passaging method; Wnt1 cells are passaged *in vivo* while TeLi cells are Wnt1-derived cells that have been established and passaged *in vitro*.

Surprisingly, this extensive and unbiased analysis did not reveal any differences in tumor infiltrating macrophage populations between either breast or pancreas tumors grown in obese and non-obese environments. This is in contrast to literature describing the influx of macrophages in obese adipose tissue (Weisberg et al, 2003). There is an assumption that increased macrophages in adipose tissue will correspond to increased macrophages in tumors. This has not been well documented and direct comparisons with the literature are difficult to make. Tumor infiltrating macrophage increases are often determined by changes in gene or protein expression and not at the single cell level (Cranford et al, 2019). The inclusion of different tumors models and their knockout counterpart analyses also make it difficult to compare macrophage abundance between obese and non-obese tumor microenvironments (Incio et al, 2016b). It is possible that tumor infiltrating macrophage differences are not in abundance and that a more macrophage focused panel might reveal macrophage phenotypic differences between the obese and non-obese setting.

DC are major antigen presenters and good targets for anti-tumor immunity therapy (Wculek et al, 2020). This mass cytometry study was not specifically designed to subset dendritic cells, but the analysis pipeline still managed to identify and characterize four DC metaclusters with distinct phenotypes. The MC21 DCs are characterized by high PD-L1 positivity indicating that this subset is T cell suppressive (Oh et al, 2020). Although the shift was small, MC21 was significantly increased in the obese E0771 group. The CD11b+ DCs found in MC16, are likely conventional cDC2 dendritic cells and are strong activators of CD4 T cells (Binnewies et al, 2019). pDC (MC10) are associated with tumor aggressiveness and poor prognosis (Wylie et al, 2019). While they were present in all groups, there was no measurable difference in abundance between the different groups, indicating that pDC do not play a major role in obesity associated cancer. The addition of CD103 to the panel would enable the identification of cDC1 subsets as well which are important for activating CD8 T cells and play a large role in anti-tumor immunity (Wylie et al, 2019).

The published research on the cancer-obesity link is sizable and growing. There are many different models and experimental designs in use. One key feature of our experimental design is the live cryopreservation of the tumor cells. This approach depletes the neutrophils, enabling a definitive identification and characterization of G-MDSC (Graham-Pole et al, 1977; Kotsakis et al, 2012). Neutrophils and G-MDSC are almost phenotypically indistinguishable by fluorescence and mass cytometry (Zhou et al, 2017; Zilio & Serafini, 2016). Since Neutrophils do not survive the freeze thaw process, CD11b+ Ly6-G+ cells in our analysis are G-MDSC and not neutrophils (Graham-Pole et al, 1977; Kotsakis et al, 2012). The multistep clustering pipeline provides useful insight into the phenotypic relatedness of these cell types. Due to differing Ly6-G expression, G-MDSC (MC19) and M-MDSC (MC18) are quite distant on the viSNE map. The strong Ly6-G staining on the G-MDSC population contributed to those cells being placed in a separate viSNE island for that metacluster. The similarity of expression for the other markers (mainly CD11b and Ly6-C) resulted in the MDSC metaclusters being very close in the hierarchical clustering. With the data available here, it would be a mistake to combine the MDSC subsets into a single metacluster. The metacluster spacing on the viSNE map makes it clear that they are distinct populations of cells, which is consistent with the literature; however a less stringent metaclustering using the dendrogram would have resulted in a single MDSC metacluster which would have been far less informative and counter to the cell spacing on the viSNE map and the literature. Although we attempted to minimize bias in this analysis pipeline, expert knowledge was still crucial for correctly subsetting these cell types. Here, we found that the G-MDSC were increased in abundance in the obese E0771 group, meaning that for the E0771 model, there are two distinct T cell suppressive cell subsets.

T cells, specifically CD8 T cells, are major players in tumor immunity. While tumor infiltrating CD8 T cells, often called TILs, are the most studied T cell type in cancer, CD4 and DN T cells are both commonly found in the tumor microenvironment. T cells play various roles in the tumor microenvironment. Here we found two phenotypically distinct and perhaps functionally distinct CD4 T cell subsets. The GITR^hi^ CD4 T cells in MC1 were also enriched for Sca-1. Both of these markers indicate an activated phenotype (Nocentini & Riccardi, 2009; Whitmire et al, 2009). GITR is also high on Tregs but there were very few CD25+ cells in this metacluster so it is unlikely that there were many Tregs in MC1. Low GITR indicates naïve or effector T cells which characterizes the second CD4 T cell metacluster, MC15. It is notable here that two distinct DN T cell populations were identified. The two DN T cell metaclusters are phenotypically distinct and far apart on the viSNE map. The Ly6-C+ DN T cells (MC5) were near the NK cells between the CD4 and CD8 T cells, while the Il-7Ra+ DN T cells (MC20) are in a separate island on the far side of the map. DN T cells are known suppressor cells and have been shown to suppress cancer cell growth in culture (Lu et al, 2019; Young et al, 2003).

CD8 T cells are the primary tumor-cytotoxic lymphocyte (Martínez-Lostao et al, 2015). CD8 T cells in cancer are often exhausted and no longer capable of cancer cell killing. Checkpoint blockade immunotherapy (CBI), such as anti-CTLA-4 and anti-PD-1/PD-L1, works to reactivate the CD8 T cells and can lead to better patient outcomes in multiple cancer types (Christofi et al, 2019). CBI only works in a small subset of patients though and it is not clear why (Zugazagoitia et al, 2016). Understanding the CD8 T cell contribution to the tumor microenvironment is key to taking full advantage of checkpoint blockade therapies. Previous reports have suggested that obese patients tend to have worse responses to traditional cancer therapies but do better with immunotherapy than non-obese patients (Wang et al, 2019). It is not clear why obese patients respond so well to checkpoint blockade immunotherapies but it may be due to the existing immune dysregulation and chronic low-grade inflammation in obese patients (Naik et al, 2019). It has been shown that patients with lower CD8 expression in the tumor microenvironment tend to have worse outcomes in response to traditional therapies (Liu et al, 2012; Mahmoud et al, 2012). It is not clear how intratumoral CD8 levels relate to CBI success or to obese patient outcome.

Here we were able to provide an in-depth characterization of multiple T cells subsets which could be invaluable for future studies in determining the therapeutic value of checkpoint blockade therapies across patients and cancer types. The two CD8 T cell subsets we found were quite similar in phenotype and both metaclusters were decreased in the obese E0771 group. This fits well with the increased abundance of two T cell suppressive cell types, G-MDSC and PD-L1+ DCs.

While diet induced obesity led to tumor immune infiltrate changes in the E0771 model, it did not have an effect across all models. Despite the systemic nature of the obese phenotype, our results suggest the interplay between obesity and tumor immune infiltrate is very cancer subtype specific. Different treatment approaches might be needed depending on the cancer subtype’s ability to alter the tumor immune infiltrate in the obese setting. It is possible that there were detectable obesity-dependent immune changes outside this 36-marker CyTOF panel. Investigating additional activation, inactivation markers, chemokine receptors, and cytokine production may shed light on those changes. But overall, our broadly defined immune panel performed well, and provided deep and robust tumor immune phenotype across models. High dimensional positional data such as imaging mass cytometry might further help unravel the obesity effect. Several recent studies have indicated that tumoral cell-cell interactions and local neighborhoods play a role in cancer severity (Jackson et al, 2020; Keren et al, 2018; Schürch et al, 2020), indicating that such spatial information add phenotypic information compared to cell frequencies obtained with suspension-based mass cytometry. The limited data from the TKO mice would suggest that the obese immune system may play a stronger role in breast cancer than in pancreatic cancer, at least at the tumor immune infiltrate level. Several studies support this concept showing that non-immune cell components contribute strongly to pancreatic cancer prognosis in obesity (Chung et al, 2020; Eibl & Rozengurt, 2019).

Here, we have presented a pipeline for analyzing large mass cytometry datasets collected across multiple time points without anchor samples. This is a powerful approach for analyzing data collected before the use of anchor samples was introduced. The multistep clustering approach allows for a reduction of biases while still allowing for expert input to guide the final metaclusters. The detailed characterization of immune infiltrate across five tumor models provides a valuable resource for planning tumor immunity studies. We found that while cell subsets were conserved across models, subset abundance was highly model specific. The inclusion of tumor immune infiltrate from obese groups, provides insight into cancer models that may or may not be relevant for studying immune infiltrate differences in obesity. We propose that the E0771 breast cancer model is a clinically relevant model for assessing immune infiltrate in obesity. G-MDSC and PD-L1+ DC suppression of T cells is clinically relevant in regard to patient care and treatment options.

## Materials and Methods

### Experimental mouse models

The Norwegian Animal Research Authority approved all animal experiments. Experiments were carried out according to the European Convention for the Protection of Vertebrates Used for Scientific Purposes. The Animal Care and Use Programs at the Faculty of Medicine, University of Bergen is accredited by AAALAC international. Male and female C57BL/6J (stock number: 000664) mice were purchased from Jackson laboratories. Mice were kept in IVC-II cages (SealsafeÒ IVC Blue Line 1284L, Tecniplast) and housed in the laboratory animal facility at the University of Bergen. Up to 6 mice were housed together and maintained under standard housing conditions at 21°C ± 0.5°C, 55% ± 5% humidity, and 12h artificial light-dark cycle. Mice were provided with food and water *ad libitium*.

At 6 weeks of age, mice in the obese cohort were placed on a high fat diet (HFD, 60% kcal from fat, 20% from protein, and 20% from carbohydrates, Research Diets, D12492) for 10 weeks, while lean mice were kept on a standard chow diet (7.5% kcal from fat, 17.5 % from proteins and 75% from carbohydrates, Special Diet Services RM1, 801151). At 16 weeks of age, mice were weighed and breast cancer cell lines were orthotopically injected in the 4th inguinal mammary fat pad for female mice, and pancreatic cancer cell lines were orthotopically injected into the lower body of the pancreas of male mice (Fig 1A). Feeding regimens were maintained throughout the experiment. Tumor growth and mouse weight were monitored over time. Cells in Phosphate Buffered Saline (PBS) were mixed 1:1 by volume with Matrigel (Corning, 356231) and injected in a total volume of 50 μl for fat pad injections and 30 μl for pancreas injections. The experiment was stopped when the first mouse showed signs of distress. At endpoint, the mice were euthanized by cervical dislocation while anaesthetized, and tumors harvested for weight measurements and downstream analysis (Fig 1A). See Table 1 for tumor model cell line details and cell injection numbers.

The C11 and UN-KC cell lines were kindly provided by Dr. Rolf Brekken and Dr. Surinder Batra, respectively. The E0771 cell line was acquired from CH3 BioSystems. The *in vivo* passaged MMTV-Wnt1 cells were kindly provided by Stein-Ove Døskeland, University of Bergen. The TeLi cell line was generated in house by *in vitro* passaging of dissociated cells from the MMTV-Wnt1 cell line injected in the mammary fat pad. The tumor was dissociated using Mouse tumor dissociation kit (Miltenyi Biotec, 130-096-730) according to manufacturer’s instructions. Dissociated MMTV-Wnt1 tumor cells were cultured *in vitro* for two months to obtain pure tumor cells now referred to as TeLi. E0771, TeLi, C11, and UN-KC cells were cultured at 37°C, 5% CO_2_ in high-glucose DMEM (Sigma, D5671) supplemented with 10% FBS (Sigma, F-7524), 100 U/mL penicillin and 100 μg/mL streptomycin (pen/strep, Sigma, P-0781) and 2 mM L-glutamine (Sigma, G-7513).

### Tumor processing and dissociation

Tumors were collected from seven different mouse cohorts. A subset of the tumors collected from those 7 cohorts became the data for the 7 batches collected on the Helios CyTOF (Table 1). Tumors were collected, weighed, minced, and incubated with collagenase II (Sigma, C6885) and DNase I (Sigma, DN25) based on the Leelatian protocol (Leelatian et al, 2017). Changes to incubation were as follows, minced tumors in RPMI-1640 (Sigma, R7388) were incubated with enzymes in capped 15 mL conical tubes in a warm water bath with periodic inversion while waiting for all tumors to be collected. Tumors were then placed in a 37°C incubator and rotated on a Ferris wheel for 1 hour. DNase was used throughout the dissociation protocol and was especially important for pancreatic tumors where free DNA was observed without the continuous use of DNase. Live cells in 10% FBS, 90% DMSO were cryopreserved with CoolCell (Sigma, CLS432001) at −80°C overnight before transfer to liquid Nitrogen.

### Determining the number of samples to barcode and number of cells to collect

Before data collection it is imperative to know the size of the target population to enable a robust analysis– particularly in heterogeneous populations. Preliminary testing on E0771 tumors suggested that the CD45+ target cells were approximately 5% of the total events collected; that held roughly true for the barcoded batches (Table 3). The following equations were used to guide the number of samples barcoded together and the time spent at the Helios for collection. The event rate was approximately 500 events per second.

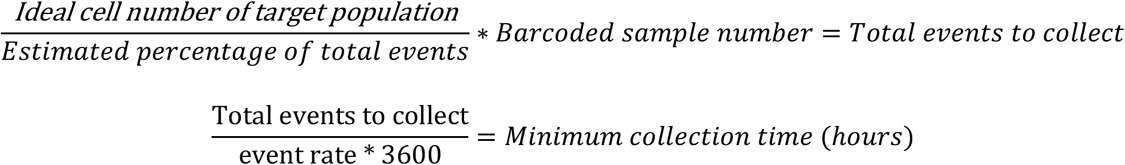

### Cell staining and running on Helios

Cells were thawed in warm water bath at 37°C. 1 mL of warm DNase supplemented media (RPMI-1640 with 10% FBS, 1% pen/step, and DNase was added to each thawed cryovial). Contents were then dumped into labeled 15 mL conical tubes containing 8 mL of warm DNase media. Cryovials were washed with 2 mL warm DNase media and added to contents to corresponding 15 mL conical tube. Cells were then rested for 5 minutes (min) followed by centrifugation at 200xg for 5 min at room temperature (RT). Cells were counted with trypan blue using a Countess automated cell counter (Invitrogen) and stained with cisplatin (Fluidigm, 201064) in the DNase media using the Fluidigm protocol. Cells were kept in warm DNase media until fixed with PFA at 1.6% final concentration. Following fixation, cells were barcoded using palladium cell barcoding kit (Fluidigm, 201060) and vendor protocol with the modification of incubating with barcodes for 45 min instead of 30 min. After barcoding, combined cells were kept in PBS + 1% BSA (Sigma A7030) and were blocked with anti-CD16/CD32 Fc block (eBoscience, 16-0161-82) and stained with a premade cocktail of antibodies shown in Table 2. Following surface staining, cells were permeabilized with 2-3 mL 100% pure cold methanol overnight at −20°C. Cells were vortexed vigorously before and after methanol addition to prevent clumping. The next day, 2 mL PBS was added to dilute methanol. Cells were spun down at 900xg and methanol/PBS mixture was pipetted off. Cells were washed with PBS again and decanting was performed as usual. Intracellular staining was performed in DPBS + 1% BSA for 30 min. After 20 min, DNase and iridium intercalator (Fluidigm, 201192B) were added. DNase was used on fixed and permed cells to reduced clogging on the Helios and cut down on visible clumping when running samples. This was especially important for the pancreatic tumors that seemed to have a high level of cell death and free DNA.

All staining was performed in capped facs tubes on a moving platform or with manual agitation to prevent cells from fully settling. Final washes were all with 3 mL followed by vortex, centrifugation at RT at 900xg, and decant: 3 PBS+1% BSA washes, 3 PBS washes, 3 milliQ washes. Cells were kept in the void volume plus 100 uL of 1X beads in MilliQ water at 4°C until ready to run. 25 μL of cells were removed at a time and added to 2 mL of diluted 1X Fluidigm EQ calibration beads in MilliQ water immediately before running on the Helios. Cell concentration was adjusted as needed so cells were running between 300 and 600 events/second. Cells were run on the Fluidigm Helios mass cytometry machine using a narrow bore injector.

A control sample was included in every batch and was used to manually check for antibody staining and machine performance. Control samples were from two different E0771 tumors that had already been phenotyped and used to test the staining panel. Controls were not used as anchor samples for batch correction because more than one control was used across the batches.

### 4T1 tumor model used for batch correction testing

Control group 4T1 breast cancer tumors from female BalbC mice from two batches were used to test the robustness of the batch correction algorithms. The 4T1 cell line was purchased from ATCC and cultured in a humidified atmosphere (37°C, 5% CO_2_). 4T1 cells were cultured in RPMI-1640 medium (Sigma) with 10% FBS, 2 mM L-glutamine, 100 U/mL penicillin and 100 μg/ml streptomycin (Sigma). MycoAlert (Lonza, LT07-318) was continuously used to confirm that the cell line was mycoplasma-free throughout the study. Female BALB/c mice (Envigo) were fed a standard chow diet. At 4-6 weeks old, 2.0 × 10^5^ 4T1 cells were mixed 1:1 with BD Matrigel Matix Growth Factor Reduced (BD Bioscience) and injected into the right mammary fat pad.

Upon sacrifice, tumors were harvested and dissociated using a mouse tumor dissociation kit (Miltenyi Biotec, 130-096-730). After dissociation erythrocytes were lysed with Red Cell Lysis Buffer (Miltenyi Biotec, 130-094-183) according to manufacturer’s protocol.

The dissociated 4T1 tumor cells were stained with cisplatin for viability with RPMI1640 (10% FBS, 0.25μM cisplatin, RT) for 5 min and fixed with PFA (1.6%, Paraformaldehyde 16% solution EM grade, Electron Microscopy Sciences, 15710) before freezing. After 10 min incubation, samples were centrifuged and supernatant was aspirated before being placed in −80°C freezer until analysis.

The panel contained 18 mass tagged antibodies with the same target as the 7 experimental batches. Many of the same antibody targets were on different channels (marked sdc in Table 4) and 1 was a different clone. A similar protocol, detailed below, was used for staining and cells were run on the same Helios CyTOF machine.

Prior to staining, samples were thawed at RT and resuspended in cell washing buffer (CWB, Dulbecco’s PBS (DPBS), (Thermo Fisher Scientific, 14040-133) with 1% BSA, 0.02% NaAzide, and 0.025% DNaseI (Sigma Aldrich, DN25-1G)). Cells were counted using Countess™ Automated Cell Counter (Invitrogen) and approximately 3 × 10^6^ cells per sample were used for barcoding. Counted cells were washed in 1X perm buffer (Fluidigm, 201057) twice, before being resuspended in 1X perm buffer (195 μl). Samples were then mixed with 5 μl barcoding solution (Fluidigm, 201060) or 1 μl (500 μM) Intercalator-103Rh (Fluidigm, 201103A) for the control and incubated for 30 min at RT. After incubation, cells were washed in CWB twice, then pooled with CWB and before washing in Maxpar® Cell Staining Buffer (CSB, Fluidigm, 201068).

Surface antibodies were diluted in CSB, while antibodies targeting intracellular proteins were diluted in CSB/perm (10% 10X perm buffer). Samples were blocked on ice for 10 min using anti-CD16/CD32 (75 μl per 3×10^6^ cells) diluted in CSB. The surface staining antibody cocktail was prepared to total 40 μl per 3 million cells. Samples were incubated with the antibody cocktail for 30 min (RT) before washing in CWB with a 10 min (RT) incubation. Samples were then washed in PBS (2 mM EDTA) and fixed with 200 μl 2% PFA per 3 million cells for 30 min (RT, in the dark). Fixed cells were then washed in CSB/perm twice and blocked with anti-CD16/CD32 in CSB/perm as described above. Samples were incubated with the intracellular staining antibody cocktail for 30 min (RT) before being washed in CWB and subsequently in 2 mM EDTA/ PBS. Cellular DNA was stained by incubating the samples in Ir191/193 Intercalator (0.33 μl 500 μM Intercalator-Ir per 2×10^6^ cells, Fluidigm, 201192B) mixed with 2% PFA (1 ml per 20×10^6^ cells) overnight (4°C). The following day, samples were centrifuged and resuspended in CWB with 10 min incubation (RT). Samples were then washed in PBS (2 mM EDTA) and kept on ice until the mass cytometer was ready. Before acquisition, an aliquot of cells was washed in MilliQ water four times (400xg, 5 min, RT). The aliquot was then resuspended in 1X EQ Four Element Calibration Beads solution (Fluidigm, 201078) to a final concentration of about 1-2×10^6^ cells per mL, and strained through a 40 μm cell strainer. The cells were then acquired on the Helios mass cytometer (Flow Cytometry Core Facility, Department of Clinical Science, University of Bergen). All centrifugation steps were performed with swing-bucket rotor at 900xg for 5 min at RT unless otherwise specified.

### Data preprocessing

Raw Helios FCS files were bulk normalized in the Fluidigm CyTOF software version 6.7.1014 using the bead normalization passport EQ-P13H2302_ver2 (Fluidigm^®^, 2018). The Fluidigm normalization tool was chosen so that all data sets could be normalized to the normalization passport. The “Original data” box was checked and the files were not concatenated at this time. The randomization was set to uniform negative distribution (UND) with linear output values, conversion compatible with FlowJo, and the default time interval normalization of 100 seconds. Beads were not removed. After normalization, the MATLAB debarcoding tool was used to simultaneously debarcode and concatenate the samples, with the exception of batch C11_1 (Zunder et al, 2015). The C11_1 raw batch files were concatenated using the Fluidigm software before debarcoding because the multiple large files were too much for the MATLAB debarcoding tool to open. Each batch was debarcoded separately with the same filter values set to minimum separation of 0.12 and a maximum Mahalanobis distance of 30.

After debarcoding, the R premessa package was used to resolve a channel naming conflict (Fig 1A) (Gherardini, 2019). The Wnt1 cohort was not stained for IRF4 so that channel was excluded for all 7 batches/cohorts. “155Gd_IRF4” was changed to “155Gd” for all FCS files and the channel was ignored during analysis. The signal was low to negative in all stained cohorts so there was minimal potential for spillover. Premessa was used more extensively in the supplemental data (9 batch dataset) to merge two very different panels so that the 18 common markers could be analyzed even if they were on different channels or had different naming conventions. The code “sdc” for shared different channel was created to denote shared antibody targets on different channels between the panels. The supplementary data also required the use of the R package cytofCore to reorder channels in the different panels (Bruggner et al, 2019).

After channel renaming with Premessa, all the newly written FCS files were uploaded to Cytobank to check for panel discrepancies and to gate for live CD45+ cells. Gating was performed manually to obtain a population of live CD45+ events (Fig 1D). The four Helios Gaussian parameters, Event_length, 140-bead channel, and Iridium (193) were gated on versus time using the cleanup strategy recommended by Fluidigm (Fluidigm^®^, 2019). Iridium intercalator was used to mark intact cells; cisplatin was used as a membrane exclusion stain to differentiate live and dead cells, Cleaved-Caspase3 (c-Cas3) was used to exclude apoptotic cells and CD45 as a pan-leukocyte marker (Fig 1D).

Gated sample files that contained fewer than 5000 live CD45+ singlets were excluded from further analysis. Because of the low viability of the pancreatic tumors, especially those from HFD, we had to remove samples from all three pancreatic cancer tumor cohorts (Fig 1C, open X means removed). Three HFD samples were removed from the C11_1 cohort, two chow and three HFD were removed from the C11_2 cohort and four HFD tumors were removed from the UN-KC cohort. No tumors were removed from the four breast cancer cohorts (Table 1).

Live CD45+ cells from the remaining 67 files (including 10 controls files) from 7 batches were exported to new FCS files and downloaded from Cytobank. The CD45+ FCS files were imported to R/RStudio for batch correction using Cydar with ncdfFlow and flowCore used as support packages (Ellis et al, 2019; Jiang et al, 2019; Lun ATL, 2017; R Core Team (2020), URL https://www.R-project.org; RStudio Team (2020), URL http://www.rstudio.com/). 35 markers were arcsinh transformed with a scale argument/cofactor of 5 before batch correction. c-Cas3 was not batch corrected since it was used only during pregating and was no longer relevant. Cydar offers three batch correction algorithms: warp, range, and quantile. All three algorithms were tested without the use of common group/anchor files. Initial batch correction algorithm testing was performed on the chow files from the 7 experimental batches and the control group files from 2 batches of 4T1 tumors from Balbc mice stained with a different panel and under different conditions to test for algorithm robustness. The 7 experimental batches were additionally batch corrected separately from the 2 testing batches using warp and range correction. Range correction was then chosen as the best algorithm for this use case and the range corrected data was used throughout the rest of the analysis. New FCS files were created binding together the original and transformed range corrected data. The 35 new corrected channels were marked with “c_” as the prefix.

### Analysis pipeline

The range corrected files were uploaded to Cytobank for analysis. viSNE was run using 26 of the corrected markers (Table 2) (Fig 3). viSNE settings were 4000 iterations on 348802 events (5206 events per FCS file) and the default settings for perplexity (30) and theta (0.5) were used. The automatic seed was 21258186. The run time was 5.58 hours and the final KL divergence was 4.75. SPADE was used for clustering with tSNE1 and tSNE2 as the clustering channels. This preserved the viSNE dimensionality reduction in the clustering process. All cells from viSNE were put into SPADE with 40 nodes/clusters chosen and the default 10% downsample. Cytobank automatic cluster gating was used to gate the events in all 40 SPADE clusters. Clusters were then manually curated and combined to reduce overclustering for G-MDSC and NK cells. The new cluster count was 37 clusters. Cell data for all experimental files was concatenated into one file for each of the 37 clusters using Premessa to concatenate the files. That data was imported into R to generate MEM scores, MEM heat map, median heat map, and hierarchical clustering (Fig 4A, EV2A). Event count numbers for clusters/metaclusters were exported for all FCS files. Excel was used to calculate abundance percentages for each metacluster out of the total CD45+ cells. In addition to exporting cluster/metacluster events counts, some manual gating was performed on the viSNE map and event counts were exported for statistical analysis (Fig 6A, B; EV3A, B). Total CD3 cells were gated on the viSNE map using CD3 marker intensity and subsequent CD3 subsets (CD4, CD8, and DN T cells) were biaxially gated using the CD4 and CD8 channels.

CITRUS was run on the E0771 data after viSNE analysis as a confirmative analysis for the metacluster statistics. CITRUS clustering was performed in Cytobank on the same 26 channels as viSNE. There were 9 files in the chow group and 8 files in the HFD group. All events were sampled with a minimum estimated cluster size of 1% (~885 events). The Significance Analysis of Microarrays (SAM) association model was used for analysis. Select significant CITRUS clusters were plotted onto the viSNE map for visualization.

### Statistics

Abundance percentages calculated in Excel were put into GraphPad Prism to perform statistics. GraphPad Prism 8.4.3 was used to plot the data and to calculate p values using multiple unpaired t tests. Consistent standard deviation was not assumed for metacluster abundance and cell subset calculations. Statistics were not applied for UN-KC and C11 due to the limited sample numbers due to low cell viability. The two batches for both C11 and E0771 were combined so that large trends would be observed and noise would be minimized.

## Data Availability

Mass cytometry data from this publication is available and can be accessed through Flow Repository (https://flowrepository.org/id/FR-FCM-Z32G).

## Acknowledgements

We thank the Flow Cytometry Core Facility, Department of Clinical Science, University of Bergen. Helios Mass Cytometer was supported by the Trond Mohn Foundation. We thank Dr. Stein-Ove Døskeland, and Dr. Surinder Batra for providing the MMTV-Wnt1, C11, and UN-KC cell lines, respectively. We thank Kanutte Huse for commenting on earlier versions of the manuscript. NH was supported by the Norwegian Cancer Society (207028) and a Trond Mohn Foundation Starter Grant.

## Author contributions

CW, and NH conceived and designed the study. CW designed and tested the antibody panel. Tumor cell injections, mouse and tumor measurements, and tumor collection was performed by LP. Tumor dissociation and single cell cryopreservation was performed by CW, HL, and PH. Cell staining and running samples on the Helios CyTOF was done by CW and HL. RB provided cell lines and study supervision. CW and NH provided direct supervision. R code was written by CW and SG. 4T1 model data was provided by SG and JL. CyTOF data analysis was performed by CW and HL. CW, SG, LP, and NH wrote the manuscript. All authors read and approved the final manuscript.

## Expanded View Figures and Legends

**Figure EV1.**
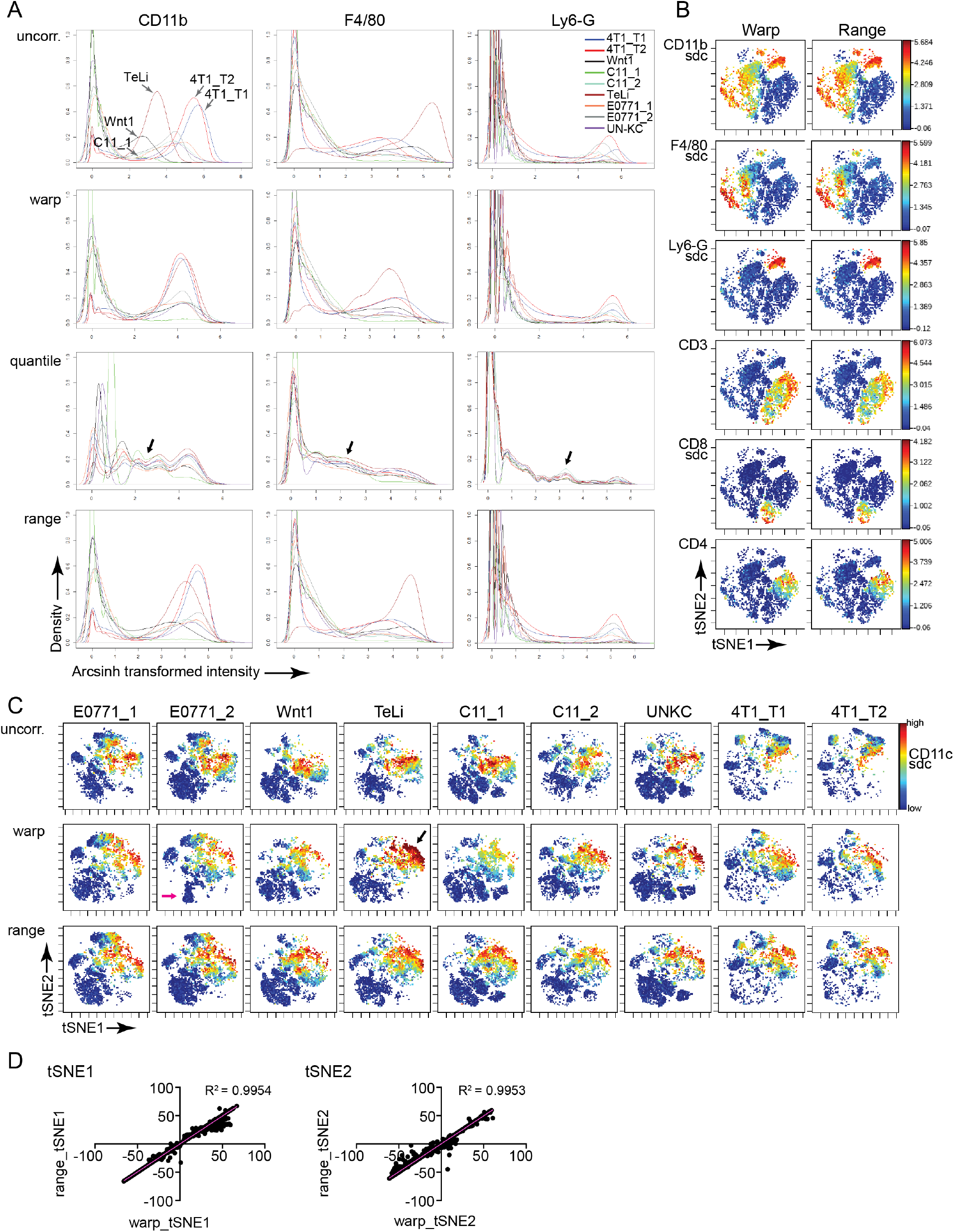
Batch correction testing with 9 testing batches using two different panels with 18 common channels including shared different channels (sdc). The 9 batches contain a total of 54 files (no HFD files) of CD45+ live cells with 5420 cells per file. A Cydar intensity distribution plots showing the third sample from all 9 batches. Intensities were plotted after arcsinh transformation for uncorrected, warp, quantile, and range corrected files. Three representative phenotyping channels are shown. Black arrows indicate the quantile density distribution that does not match the original uncorrected data. B Representative viSNE marker heat plots from 1 sample (E0771_1-C1) from a viSNE run where the warp and range corrected files for all 9 batches were run together. Range and warp plots are shown on the same scale; intensity heat scales vary between markers. C Three separate viSNE runs using the same seed (1794942912) were performed for each: uncorrected, warp corrected, and range corrected datasets. Black and pink arrows indicate warping artifacts. D Linear regression with R squared values comparing warp and range tSNE coordinates from data in panel B.

**Figure EV2.**
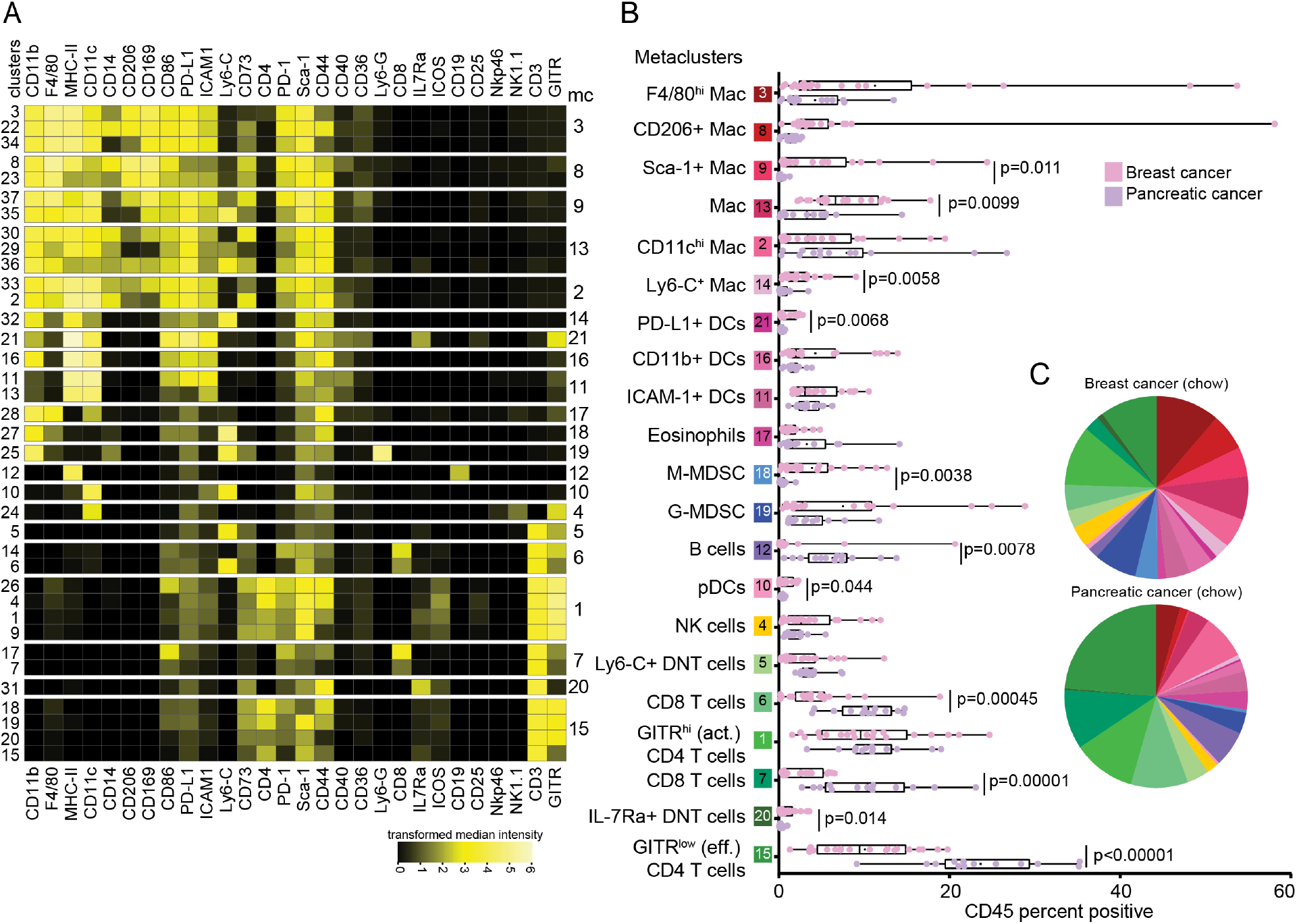
Immune infiltrate metacluster abundance between breast and pancreatic cancer. A Heat map of arcsinh transformed batch corrected median marker intensity for the 37 clusters (left labels) and 28 markers (bottom labels). The 21 metaclusters are indicated by row spacing and numbers to the right. Row order is the same as in Figure 4A. B Min to max box and whiskers plots with all points showing, median is the vertical line, dot is the mean. The plots show metacluster abundance from the combined breast cancer and pancreatic cancer non-obese murine tumors. Significant p values are displayed on the plot. C Pie charts showing the relative mean abundance of the immune metaclusters between pancreatic and breast cancer tumor models (data from chow-fed non-obese murine tumors only).

**Figure EV3.**
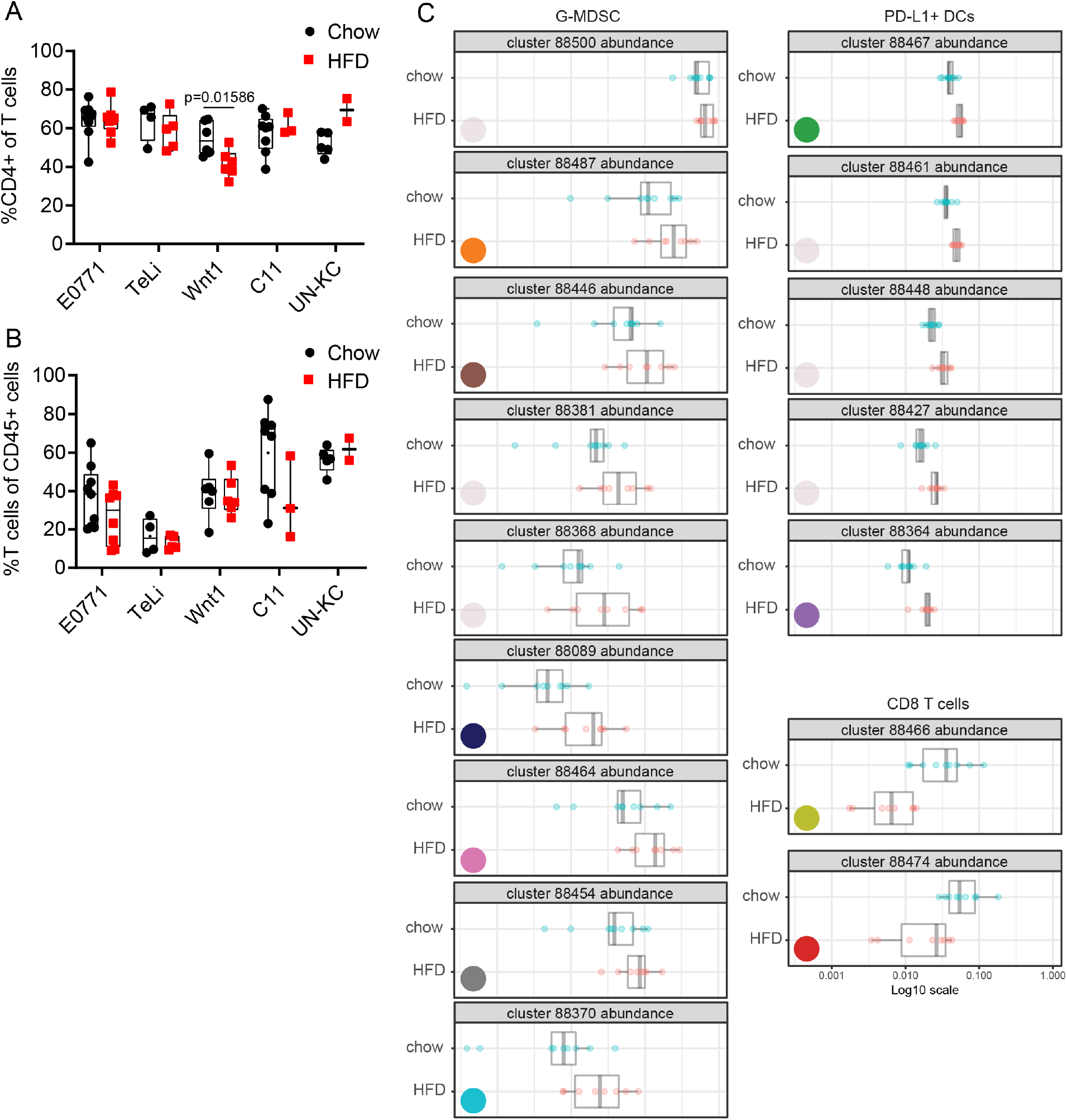
T cell analysis and CITRUS significant cluster abundance. A Percent of CD4 T cells out of total T cells for all tumor models. B Percent of T cells out of total CD45+ immune infiltrating cells for all tumor models. C Abundance plots for significant CITRUS clusters from E0771 SAM CITRUS analysis. Color-coding corresponds to Figure 6C, D.

